# Compulsive coping behaviour, developed predominantly by sign-trackers, is exacerbated by chronic atomoxetine

**DOI:** 10.1101/2024.10.08.617254

**Authors:** Chloe S. Chernoff, Aude Belin-Rauscent, Mickaël Puaud, Sebastiano A. Torrisi, Maxime Fouyssac, Bence Németh, Charlotte (Zhixuan) Yu, Alejandro Higuera-Matas, Sue Jones, David Belin

## Abstract

**Background:** Loss of control over coping strategies can result in the development of impulsive/compulsive spectrum disorders (ICSDs) such as obsessive-compulsive disorder or trichotillomania. Rats, like humans, show individual differences in their tendency to engage in and maintain control over coping behaviours. While most rats exposed to a schedule-induced polydipsia (SIP) procedure develop controlled, moderate, polydipsic coping, some vulnerable individuals engage in excessive, compulsive drinking, or *hyper*dipsia. The development of hyperdipsia depends in part on noradrenergic mechanisms, as it is prevented by the noradrenaline reuptake inhibitor, atomoxetine in highly impulsive vulnerable rats. However, whether noradrenergic mechanisms also underlie the expression of well-established hyperdipsia, or if other traits, such as the ICSD-relevant sign-tracking, confer vulnerability to its development, are unknown.

**Methods:** In two longitudinal studies in male Sprague-Dawley rats, we investigated whether well-established hyperdipsia is influenced by atomoxetine and whether its development is predicted by sign-tracking.

**Results:** Sign-tracking was associated with faster acquisition of SIP and the development of high or compulsive levels of SIP. Chronic atomoxetine both exacerbated hyperdipsia and increased the mRNA levels of the markers of cellular activity and plasticity c-fos and zif268, across the dorsal striatum, as revealed by qPCR assays. Atomoxetine also altered the transcriptomic landscape of the nucleus accumbens shell and the pattern of cFos and zif268 expression in the amygdalo-striatal system.

**Conclusions:** These results provide new insight into the biobehavioural basis of compulsive behaviours, revealing a differential noradrenergic control of the development and expression of compulsive coping, the latter involving recruitment of distinct striatal processes.

## Introduction

The ability to adaptively cope with stress and regulate negative emotions is paramount for resilience and wellbeing (1–3). However, some coping strategies become excessive and rigid, characteristic of psychiatric disorders such as mood/anxiety disorders (4) and impulsive/compulsive spectrum disorders (ICSDs) (5, 6), including substance use disorders (7).

Adjunctive behaviours are a form of displacement behaviour expressed by many species as a means of coping with stress (8). An example of such coping behaviour is schedule-induced polydipsia (SIP), wherein food-deprived rats drink supra-homeostatic volumes of water in response to the stress induced by predictable, intermittent food delivery (9, 10). SIP is anxiolytic, as it reduces plasma corticosterone levels (10–12) and anxiety-like behaviour (13, 14), which are both increased by periodic food delivery in food-restricted individuals (14, 15).

While most rats maintain a stable, controlled level of SIP, some lose control over their coping response and develop maladaptive, excessive SIP (16–18). This *hyper*dipsia is compulsive in nature as it results in animals expending energy they don’t have (as they are in a caloric deficit state) to consume, warm, and excrete water they do not need (19, 20). While the acquisition of such a coping response depends on the mesolimbic dopaminergic system (21), well-established compulsive hyperdipsia is controlled by the dopamine-dependent anterior dorsolateral striatum (aDLS) habit system (13). This shift in the neural locus of control is analogous to that accompanying compulsive drug-seeking habits and OCD-like repetitive behaviours (22, 23)

The biobehavioural factors contributing to the individual tendency to develop compulsive coping behaviours are not yet well understood. Motor impulsivity, i.e., the tendency to act prematurely, has been associated with the individual vulnerability to develop compulsive cocaine taking (24), compulsive cocaine-seeking habits (25) and hyperdipsia (26), thereby representing an endophenotype of vulnerability to compulsive behaviours. However, impulsivity explains only a fraction of the tendency to develop compulsive behaviours. Trait sign-tracking, i.e., the propensity to approach reward-predictive Pavlovian conditioned stimuli (CSs) (26–28), which is associated with the tendency to make risky choices (29) and to develop compulsive heroin seeking (Fouyssac et al., unpublished data) may contribute to SIP or its compulsive manifestation through Pavlovian processes (30–34).

At the neurochemical level, dopaminergic mechanisms in the nucleus accumbens (NAc) underpin sign-tracking, motor impulsivity and the development of SIP (35–39). The neurochemical mechanisms underlying hyperdipsia, however, appear to be distinct (11, 26, 40). Thus the development of hyperdipsia is associated with transcriptional alterations in the noradrenergic locus coeruleus (LC) (11) and is prevented by the selective noradrenaline reuptake inhibitor atomoxetine, which also decreases impulsivity in hyperdipsia-vulnerable highly impulsive rats (26) characterised by heightened NAc noradrenaline (11, 41–43). It remains unclear whether similar noradrenergic mechanisms orchestrate the maintenance of well- established, aDLS dopamine-dependent hyperdipsia or if they mediate the potential link between NAc-dependent behaviours, such as sign-tracking, and the development of compulsion.

In two longitudinal studies using male Sprague-Dawley rats, we investigated whether sign- tracking confers vulnerability to the development of hyperdipsia and whether well-established hyperdipsia or sign-tracking is influenced by atomoxetine. Using post-mortem molecular biology assays, we also characterised the influence of atomoxetine on hyperdipsia-related cellular activity within the corticostriatal circuitry and expression patterns of candidate genes in the NAc shell (NAcS).

## Methods and Materials

### Subjects

One hundred and eight 10-week-old adult male Sprague-Dawley rats, housed as described in the **Supplementary Online Materials (SOM)**, were used in this study. All procedures were conducted under the project license 70/8072 held by David Belin in accordance with the requirements of the UK Animals (Scientific Procedures) Act 1986, amendment 2012.

### Experimental design

Experiment 1 (n=48) aimed to examine the relationship between sign-tracking and the propensity to develop polydipsia or its compulsive manifestation, hyperdipsia. Rats were first assessed for their sign-tracking or goal-tracking tendency in an autoshaping task prior to being exposed to a SIP procedure, as described in the **SOM** and **Figure 1A-B**. Experiment 1 also aimed to assess the influence of acute manipulation of noradrenergic signaling on goal-tracking, sign-tracking, and SIP/hyperdipsia via acute atomoxetine challenge following training on both autoshaping and SIP procedures (**Figure 1A-B**). Experiment 2 investigated, in a separate cohort (n=60), the behavioural, neural and cellular effects of chronic atomoxetine treatment on well-established, habitual aDLS dopamine-dependent hyperdipsia (13). The same treatment was previously shown to prevent the development of hyperdipsia in vulnerable, highly impulsive rats (26) (**Figure 1C**).

**Figure 1.**
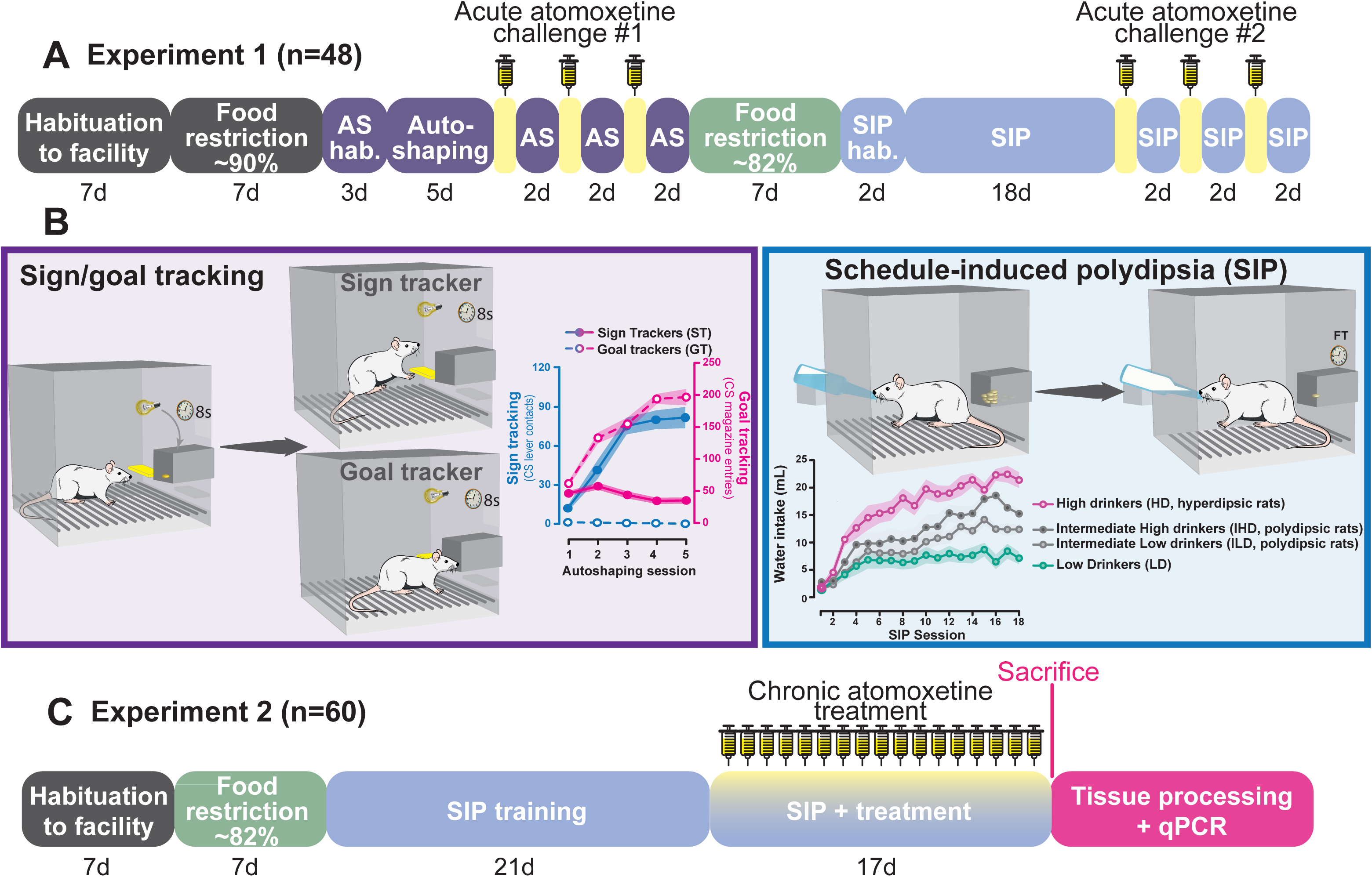
Timeline of the experiments and behavioural tasks. **A)** Following 7 days of habituation after arrival at the facility, rats were food restricted to 85-90% of their theoretical free-feeding weight. Across the three consecutive days prior to autoshaping training **(B-left)**, rats underwent habituation to sugar pellets and the autoshaping chamber, and basic magazine training. Next, rats were trained on the autoshaping task (AS) for five sessions, after which the atomoxetine challenge began according to a counterbalanced Latin square design (see SOM for details). Rats were then further food restricted to 80-85% of their theoretical free-feeding weight, and habituated to the SIP chamber and the fixed time 60-second (FT-60) food pellet delivery schedule **(B-right)**. SIP training was then performed for 18 60-min sessions. The second atomoxetine challenge occurred according to the schedule as for the previous Latin square. **B)** During the autoshaping, Pavlovian conditioned approach task, rats were exposed daily to 25 presentations of the lever and cue light above [ie. The compound conditioned stimulus (CS)] for 8 seconds, after which the lever was retracted, the stimulus light turned off, and one food pellet was delivered into the magazine. CSs-US pairs were presented according to a 90-second variable interval schedule. Sign-tracking was quantified by the number of lever presses made during the CS presentation, as an index of approach toward and interaction with either the light or the lever of the CS. Goal-tracking was measured by the number of nose pokes made into the food magazine during the CS presentation. During SIP, the majority of the population develops low or high levels of polydipsic drinking, an adaptive coping response while some animals do not develop a coping response with water and others instead develop hyperdipsia, an excessive and compulsive coping response. **C)** In Experiment 2, following acclimatization to the facility, rats were food-restricted to 80-85% of theoretical free-feeding weight. During the last two days of food restriction, rats underwent habituation and magazine training as in Experiment 1. Rats were then performed SIP for 21 days, during which high, high intermediate, low intermediate (ie. Adaptive) and low adjunctive drinkers were identified. Equal numbers of rats from each drinking group then received intraperitoneal (i.p.) injections of atomoxetine (1mg/kg) or vehicle 30 mins before each daily SIP session for 17 days. Forty-five minutes after the final SIP session, rats were sacrificed and their brains were harvested and flash frozen for subsequent qPCR analyses.

### Behavioural training and classification

The autoshaping task was conducted over 5 sessions as described previously (44) and detailed in the **SOM**. Rats were identified as sign-trackers (STs), intermediate performers (INT), or goal-trackers (GTs) using a K-means cluster analysis of sign-tracking and goal-tracking behaviours across the last three autoshaping sessions.

SIP training was carried out over 18-21 1-hour daily sessions under a fixed time schedule of 45 mg pellet delivery, as described previously (26) and detailed in the **SOM**.

Rats displaying high or low levels of polydipsic behaviour (high intermediate adjunctive drinkers, IHD, and low intermediate adjunctive drinkers, ILD, respectively), and those which demonstrated excessive, hyperdipsic, coping (high drinkers, HD) or very low, if any, SIP (low drinkers, LD), were identified based on a quartile split of water consumption over the last three SIP sessions (as previously described (11, 26) and detailed in the **SOM**).

### Drugs

Atomoxetine hydrochloride (ATO, Sequoia Research Products, UK) was dissolved in sterile 0.9% saline vehicle. Atomoxetine (acute: 1.0, 3.0mg/kg; chronic: 1.0mg/kg; all at 1.0mL/kg) and vehicle were administered intraperitoneally (i.p) 30 minutes before behavioural sessions (26, 45, 46) as detailed in the **SOM**.

### Molecular biology analyses

Forty-five minutes after the final SIP session, rats in Experiment 2 were anaesthetised and decapitated. Brains were rapidly extracted and flash frozen in -40°C isopentane. Brains were processed as described in the **SOM** and cDNA libraries were synthesised from RNAs extracted from brain regions involved in impulsivity, compulsion and the associated transition from goal-directed to habitual control over behaviour: anterior and posterior insular cortex (aIC, pIC), nucleus accumbens shell and core (NAcS, NAcC), anterior dorsolateral striatum (aDLS), posterior dorsomedial striatum (pDMS), and basolateral and central amygdala (BLA, CeA) (13, 14, 23, 47, 48). Relative mRNA levels were determined in duplicate runs using qPCR (see 49 and **SOM** for more details). Relative gene expression levels from the brain regions of interest were used to: 1) compare SIP-related cellular activity patterns between adjunctive drinking (HD, LD) and treatment (atomoxetine, vehicle) groups using an immediate early gene (IEG) hotspot analysis (50); 2) assess relationships between levels of adjunctive drinking behaviour and IEG activation across striatal and limbic regions as a proxy of adjunctive drinking-related brain activation; and 3) conduct an exploratory network analysis to examine transcription patterns within IEG, catecholamine, and endogenous opioid systems of the NAcS – the only striatal subregion receiving notable noradrenergic innervation (11, 51, 52). Genes of interest and respective primers are listed in **Table S1**.

### Data and statistical analyses

Data, presented as mean ± SEM and/or individual data points, were analysed using SPSS 28.0.1 (IBM Corporation, USA) as detailed in the **SOM**.

## Results

### Experiment 1

#### Sign tracking confers a tendency to engage in high levels of coping behaviour and vulnerability to develop hyperdipsia

In the autoshaping task, distinct groups were revealed by k-means clustering based on ST and GT approach behaviours (**Figure 2A**). STs made more CS-concurrent lever contacts than GTs (**Figure 2B**), which instead entered the food magazine more than the former during CS presentation (**Figure 2B)**. After more than three weeks of exposure to a SIP procedure, these rats were shown to engage in different degrees of coping behaviour. HD rats exhibited compulsive coping, or hyperdipsia, with much higher levels of SIP than IHD and ILD rats, which demonstrated high or low levels of adaptive coping behaviour, and LD rats, which did not readily develop SIP with water, as previously reported (13, 18) (**Figure 2C**). HD rats quickly lost control over their coping responses, since they drank more than IHD, LHD and LD rats from the 8^th^, 6^th^, and 3^rd^ SIP session onward, respectively (**Figure 2D**). IHD and ILD rats differed from LDs in their coping response from sessions 12 and 13, respectively, while differences in SIP levels between them emerged after the 14^th^ session (**Figure 2D**).

**Figure 2.**
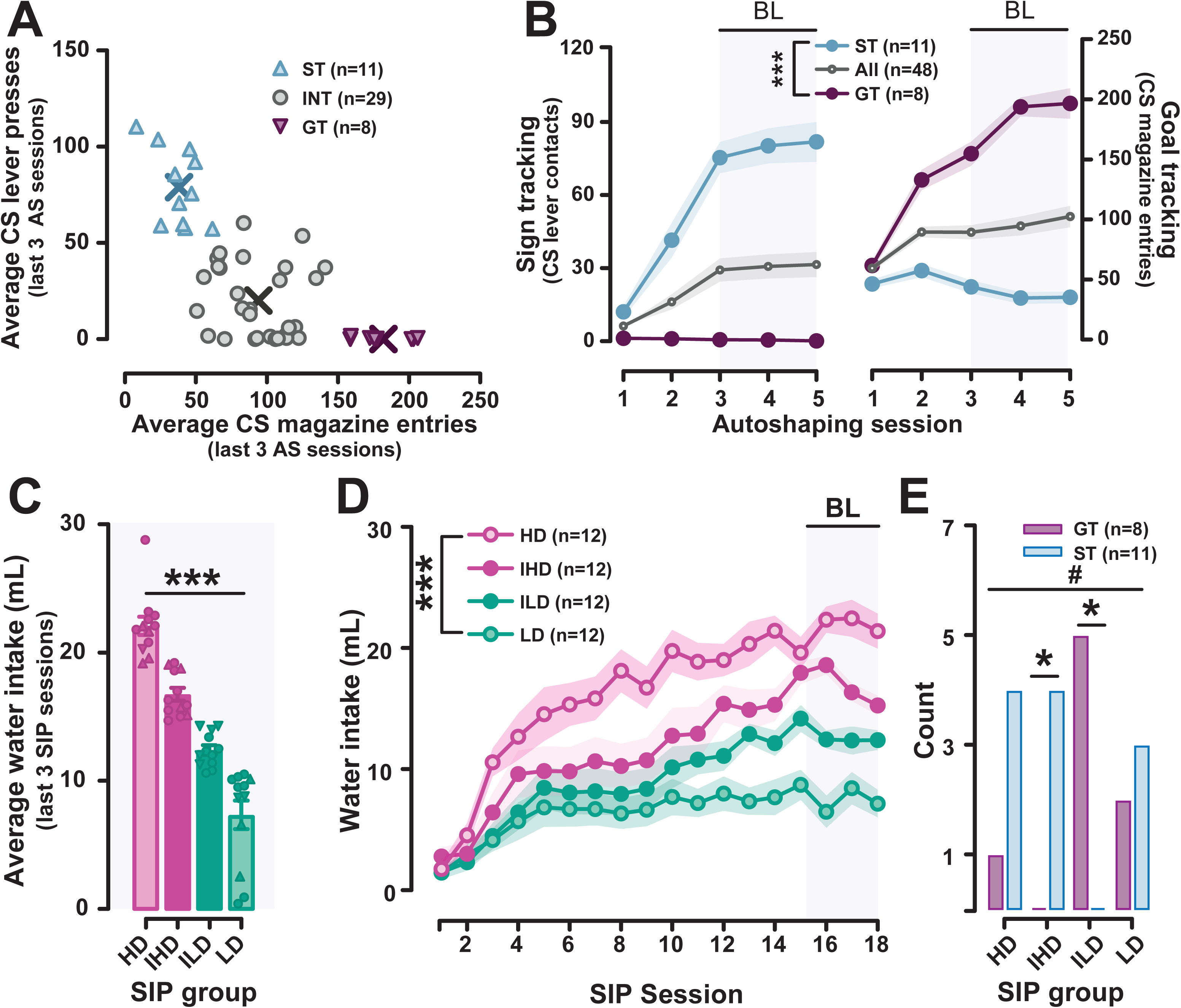
Characterization of individual differences in sign-tracking, goal-tracking and coping behaviour. **A)** A k-means clustering algorithm was used to categorize rats as sign trackers (STs), goal trackers (GTs), or intermediate performers (INT) based on average lever presses and magazine entries made during the CS-presentation across the last three autoshaping training sessions. The “X”s represent cluster centres. **B)** As expected, STs and GTs demonstrated greater responding either on the lever [session x group: F_8,180_ = 17.240, p < 0.001, η^2^ = 0.434; GTs vs STs- all sessions: Sidak ps ≤ 0.003, ds ≥ 0.504] or at the food magazine [session x group: F_8,180_ = 23.766, p < 0.001, η^2^ = 0.514; GTs vs STs- session 1: Sidak p = 0.094; sessions 2-5: Sidak ps ≤ 0.001, ds ≥ 1.165], respectively, during the CS presentation. **C)** Rats were defined as high (HD), high intermediate (IHD), low intermediate (ILD), or low (LD) drinkers based on a quartile split of average water consumption across the last 3 days of SIP training. **D)** Differences between SIP groups emerged at various points in acquisition, such that by the end of SIP training, groups exhibited distinct levels of water consumption across the last three SIP training sessions (BL) [session x group: F_51,782_ = 3.402, p < 0.001, η^2^ = 0.188; HD vs ILD- sessions 1-5: Sidak ps ≥ 0.087; sessions 6-18: Sidak ps ≤ 0.036, ds ≥ 0.416; HD vs IHD- sessions 1-7: Sidak ps ≥ 0.188; sessions 8-18: Sidak ps ≤ 0.012, ds ≥ 0.475; HD vs LD- sessions 1-2: Sidak ps ≥ 0.770; sessions 3-18: Sidak ps ≤ 0.042, ds ≥ 0.408; [IHD vs LD- sessions 1-11: Sidak ps ≥ 0.085; sessions 12-18: Sidak ps ≤ 0.003, ds ≥ 0.545; ILD v LD- sessions 1-12: Sidak ps ≥ 0.534; sessions 13-18: Sidak ps ≤ 0.054, ds ≥ 0.393; IHD vs ILD- sessions 1-14: Sidak ps ≥ 0.152; sessions 15-18 ps ≤ 0.037, ds ≥ 0.393]**. E)** STs and GTs were not evenly distributed across SIP groups [χ^2^_3,n=19_ = 10.795, p = 0.013]. ST rats were 8 times more likely to develop high or excessive SIP than GT rats [HD/IHD vs ILD/LD: Chi^2^= 7.59, p < 0.01] with but one GT rat present in the HD and IHD groups as compared to 8 ST rats. In contrast, there were fewer STs vs GTs in the ILD group [ILD- GT vs ST: z = -3.055, p = 0.002]. As a matter of fact, there were significantly more GT rats in the ILD group than any other SIP group [GTs- ILD vs HD: z = - 2.582, p = 0.010; ILD vs IHD: z = -3.000, p = 0.003; ILD vs LD: z = -2.070, p = 0.038; all other |zs|≤ 1.920, ps ≥ 0.055] while there were only STs rats in the IHD group [ST vs GT: z = -1.920, p = 0.055]. Data are presented as A) individual datapoints representing each subject, B-D) group means ± SEM, or E) number of subjects per group. Symbols in C) indicate autoshaping group (GT = inverted triangle; ST = triangle). **\*\*\***significant Sidak-corrected group differences between all groups. **\***and line: p ≤ 0.05 according to a z-test of independent proportions.

There was a profound enrichment of ST rats in the HD and IHD groups. In fact, ST rats were 8 times more likely to develop high or excessive SIP than GT rats, with the HD and IHD groups containing but one GT rat. In contrast, the ILD group contained no ST rats (**Figure 2E**), and there were more GT rats in the ILD group than in any other SIP group (**Figure 2E**).

GTs and STs also differed in the speed at which they acquired SIP. STs drank significantly more water than GTs during early SIP (d1-9) (**Figure 3A**). As SIP became more established, GTs eventually reached comparable levels of SIP to STs so that, in spite of their differences in acquisition tendency and vulnerability to develop high or compulsive-like levels of SIP as described above, the difference in in the level of adjunctive drinking of GT and ST rats during later training sessions (d10-18) did not reach statistical significance (**Figure 3A**). Rats which developed hyperdipsia (HD) did not differ from non-polydipsic LD rats in goal-tracking magazine entries nor in sign-tracking lever presses (**Figure 3B**). However, ILD rats had made fewer sign-tracking lever contacts and more goal-tracking magazine entries than both IHD and HD rats (**Figure 3B**), thereby indicating that Pavlovian conditioned approach tendencies are expressed to the lowest degree in the group of rats which adopted a controlled adaptive coping strategy. Goal-tracking and sign-tracking of adaptive ILD rats did not significantly differ from that of the LD group (**Figure 3B**).

**Figure 3.**
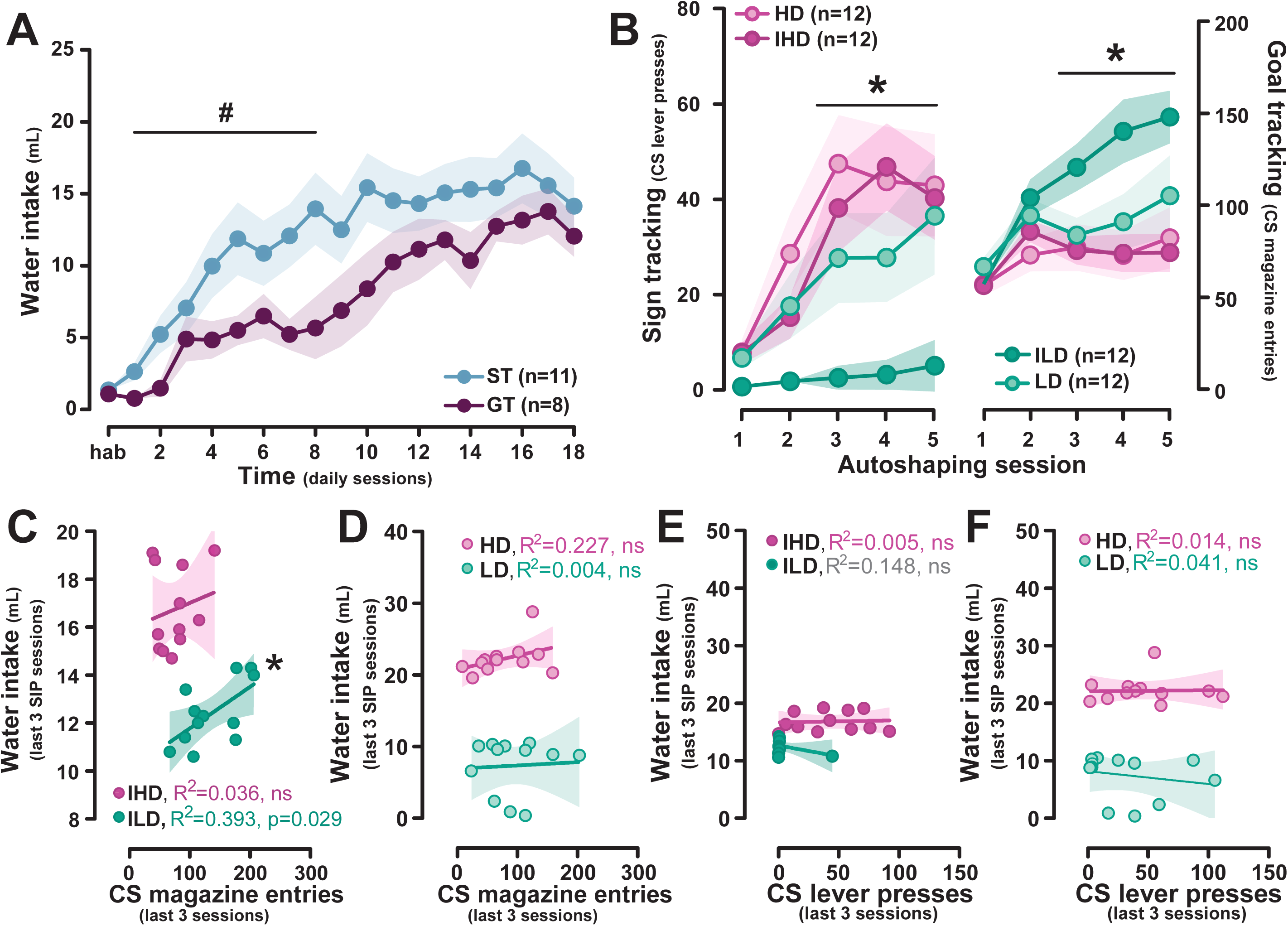
Sign-tracking is associated with faster development of SIP, while goal-tracking predicts controlled adjunctive coping. **A)** STs acquired polydipsic coping under SIP earlier than GTs [group x block x session: F_8,136_ = 2.722, p=0.008, η^2^ = 0.138; early block- session x group: F_8,136_ = 2.375, p=0.020, η^2^ = 0.123; GTs vs STs- sessions 1,2,7,8: Sidak ps ≤ 0.033, ds ≥ 0.335; sessions 3-6: Sidak ps ≥ 0.057, ds ≤ 0.294]. However, as SIP training progressed, GTs reached comparable levels of SIP to STs [late block- session x group interaction: F_8,136_ = 1.330, p = 0.234]. **B)** Autoshaping behaviour did not differ between rats that developed hyperdipsia (HD) and those which did not develop adjunctive drinking (LD) [CS magazine entries- session x group: F_12,176_ = 3.968, p < 0.001, η^2^ = 0.213; all sessions Sidak ps ≥ 0.413; CS lever presses- session x group: F_12,176_ = 2.949, p = 0.007, η^2^ = 0.167; HD vs LD: all sessions Sidak ps ≥ 0.485]. However, goal-tracking and sign-tracking differed between rats that expressed more intermediate levels of polydipsia, whereby adaptive low intermediated drinkers (ILD) exhibited a more goal-tracker-like phenotype, having made fewer CS lever presses [ILD vs IHD- sessions 3-5: Sidak ps ≤ 0.027, ds ≥ 0.431; sessions 1-2: Sidak ps ≥ 0.515; ILD vs HD- sessions 2-5: Sidak ps ≤ 0.048, ds ≥ 0.400; session 1: Sidak p = 0.089] and more CS magazine entries [ILD vs IHD- sessions 3-5: Sidak ps ≤ 0.030, ds ≥ 0.426; session 1-2: Sidak ps ≥ 0.598; ILD vs HD- sessions 3-5: Sidak ps ≤ 0.040, ds ≥ 0.410; sessions 1-2: Sidak ps ≥ 0.080] than both high intermediate drinkers (IHD) and compulsive HDs. Autoshaping behaviour between ILD and LD groups did not significantly differ [CS lever presses: all sessions Sidak ps ≥ 0.146; CS magazine entries: all sessions Sidak ps ≥ 0.110]. **C-D)** Simple linear regression indicated that both approach tendencies together did not predict water consumption under SIP [HD: F_2,9_ = 1.980, p = 0.194; IHD: F_2,9_ = 0.383, p = 0.693; ILD: F_2,9_ = 3.052, p = 0.097; LD: F_2,9_ = 0.297, p = 0.769], yet goal-tracking magazine entries alone predicted water consumption, albeit only in the ILD group [ILD: F_1,10_ = 6.462, p = 0.029; HD: F_1,10_ = 2.943, p = 0.117; IHD: F_1,10_ = 0.371, p = 0.556; LD: F_1,10_ = 0.040, p = 0.845, R^2^ = 0.393; average ILD water consumption = 10.032 + 0.017 x (average CS magazine entries)]. **E-F)** Sign-tracking lever presses, however, did not predict SIP water consumption in any group [all R^2^ s ≤ 0.148, ps ≥ 0.217]. For A-B), data are presented as group means ± SEM. Individual values fit with the linear regression line ± a 95% confidence interval are plotted in C-F). **#**p<0.05 Sidak-corrected group comparison for the indicated block. *line: p<0.05 Sidak-correct group comparison between ILD group and both IHD and HD groups. *alone: significant linear regression.

A multivariate linear regression model that was used to estimate the extent to which the level of adjunctive drinking could be predicted by both goal-tracking and sign-tracking did not yield any statistically significant result in any of the SIP phenotypes. However, simple linear regression revealed that goal-tracking magazine entries alone predicted the level of adjunctive drinking in the ILD group (**Figure 3C-D**). Thus, while goal-tracking accounted for 39.3% of the variance in adjunctive drinking in ILD rats (**Figure 3C**), it did not predict their baseline, regulatory drinking [R^2^ = 0.003, p = 0.863] (see **SOM**), and sign-tracking alone predicted neither the level of regulatory drinking (see **SOM**) [R^2^ = 0, p = 0.962] nor that of adjunctive drinking for any SIP group [all R^2^s ≤ 0.148, ps ≥ 0.217] (**Figure 3E-F**).

#### Acute atomoxetine treatment does not influence Pavlovian approach or polydipsic coping

Acute atomoxetine administration did not influence sign-tracking (**Figure 4A**) but it resulted in a slight group-dependent decrease in goal tracking (**Figure 4B**). The high dose of atomoxetine tended to decrease magazine entries in GTs (**Figure 4B**), while the low dose reduced this behaviour in ILD rats (**Figure 4B**), characterised by a particularly high level of goal-tracking (see **Figure 3D**). Consequently, the effect of atomoxetine on goal-tracking in ILD rats no longer reached statistical significance when baseline magazine entries were used as a covariate in the above analysis (**Figure 4B**).

**Figure 4.**
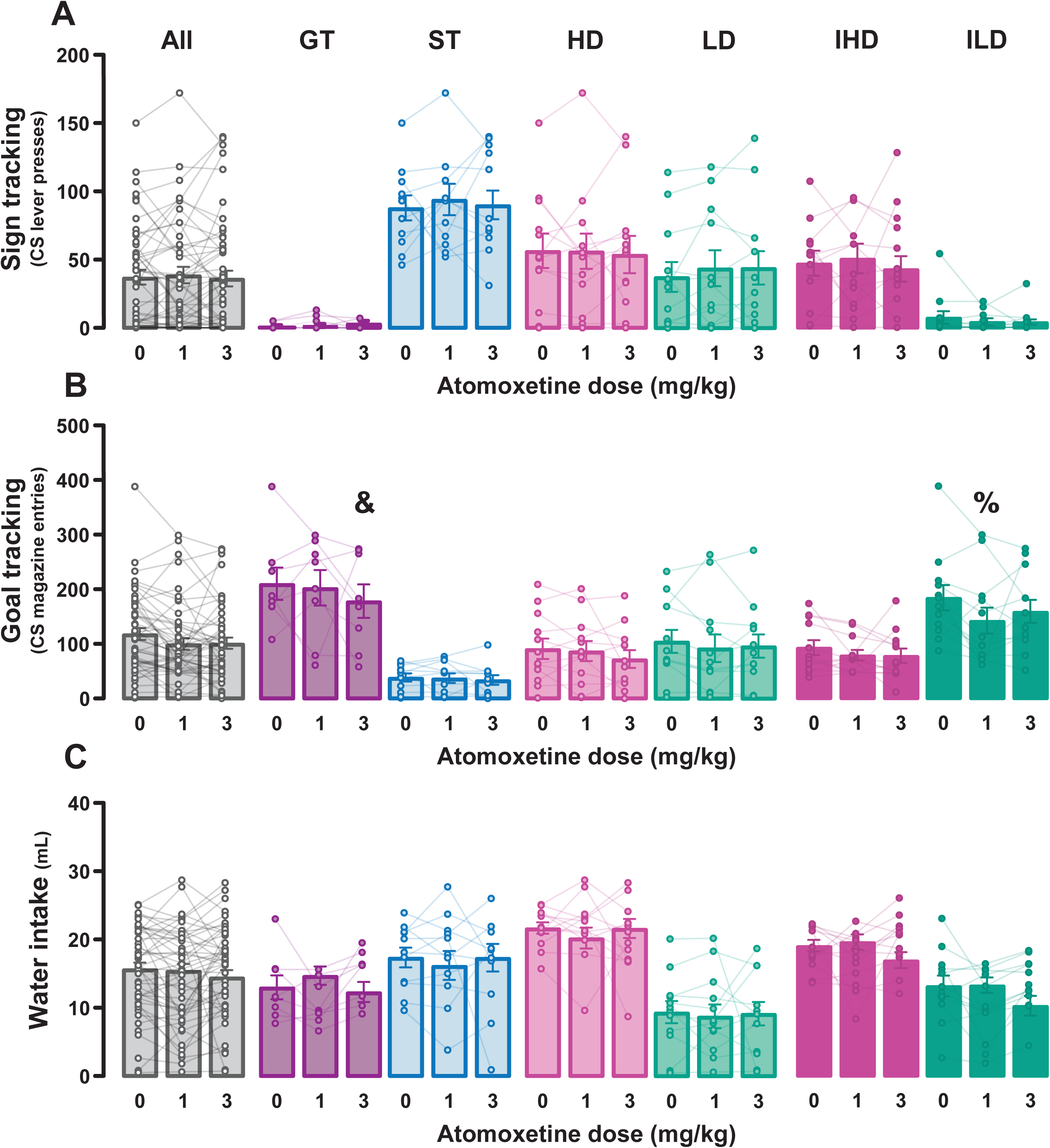
Acute atomoxetine administration has negligible effects on autoshaping and schedule-induced polydipsia. **A)** Atomoxetine did not influence sign-tracking in any group [Fs ≤ 0.237, ps ≥ 0.880], yet, at the high dose, it produced a trend-level reduction in goal-tracking in GTs only [dose x group: F_4,76_ = 3.160, p = 0.019, η^2^ = 0.143; GTs- 3.0 vs vehicle: Sidak p = 0.055, d = 0.354]. **B)** The low dose of atomoxetine reduced goal-tracking behaviour selectively in ILD rats [ILD- 1.0 vs vehicle: Sidak p = 0.002, d = 0.536], yet this effect appeared to be driven by the high rate of CS magazine responding in these rats, as this effect did not survive an analysis of covariance controlling for baseline CS magazine entries [group x dose: F_6,74_ = 2.240, p = 0.060; ILD- 1.0 vs vehicle: Sidak p = 0.372]. **C)** Acute atomoxetine did not affect well-established polydipsic responding in any group [dose: F_2,76_ = 2.911, p = 0.061, η^2^ = 0.071; dose x group: Fs ≤ 2.301, ps ≥ 0.096]. Data are presented as group means ± SEM. **&**Sidak p<0.056 compared to vehicle. **%**Sidak p<0.05 versus vehicle, but p>0.05 with baseline CS magazine entries as a covariate.

Acute administration of atomoxetine did not significantly affect either the well-established expression of adaptive or compulsive coping behaviour (**Figure 4C**) in marked contrast with the effect of a chronic administration of the drug on either the development of hyperdipsia (see 26) or its expression, as shown below.

### Experiment 2

#### Chronic atomoxetine treatment exacerbates compulsive coping behaviour

As in Experiment 1, individual differences in adjunctive coping behaviour appeared after three weeks of exposure to a SIP procedure in a separate cohort of rats (**Figure 5A&B**). HD rats in Experiment 2 lost control over their coping behaviour, showing levels of SIP significantly much higher than those shown by LD (**Figure 5C&D**), ILD, and IHD rats (**Figure 5E**) from session 2, 11, and 18, respectively. IHD rats demonstrated higher levels of SIP than LD (**Figure 5C&D**) and ILD (**Figure 5E&F**) rats from day 9 and 12 onward. The difference in water consumption between ILD and LD rats emerged after session 19 (**Figure 5E&F**).

**Figure 5.**
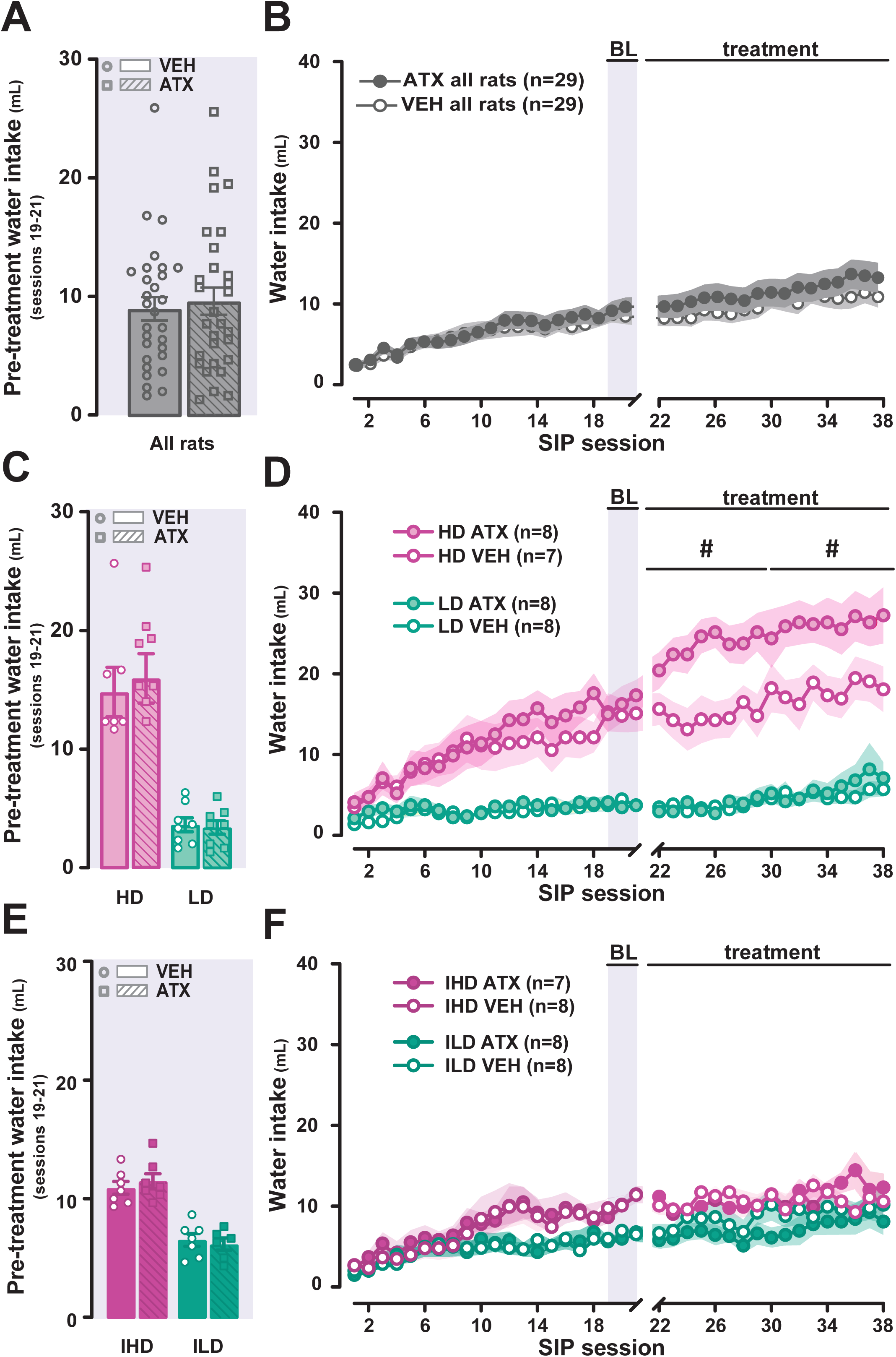
Chronic atomoxetine treatment selectively exacerbates well-established hyperdipsia in compulsive-like, high-drinking rats. **A-B)** Distinct SIP phenotypes emerged across 21 days of SIP training [session x group: F_60,980_ = 4.952, p < 0.001, η^2^ = 0.408; HD vs LD- session 1: p = 0.066; sessions 2-21: ps ≤ 0.054, ds ≥ 0.392; HD vs ILD- sessions 1-10: ps ≥ 0.081; sessions 11-21: ps ≤ 0.029, ds ≥ 0.380; HD vs IHD- sessions 1-17: ps ≥ 0.173; sessions 18-21: ps ≤ 0.043, ds ≥ 0.300; IHD vs LD- sessions 1-8: p ≥ 0.073; sessions 9-21: ps ≤ 0.007, ds ≥ 0.445; IHD vs ILD- sessions 1-11: Sidak ps ≥ 0.138; sessions 12-21: ps ≤ 0.044, ds ≥ 0.403; ILD vs LD- sessions 1-19: Sidak ps ≥ 0.140; sessions 20-21: ps ≤ 0.034, ds ≥ 0.382]. **A,C,E)** Groups that went on to receive chronic atomoxetine or vehicle treatment were randomly assigned and matched for baseline water consumption [treatment group: F_1,50_ = 0.129, p = 0.721]. **D)** Hyperdipsia displayed by HD rats was selectively exacerbated by chronic atomoxetine throughout the entirety of treatment [group x treatment x block: F_9,147_ = 3.656, p = 0.003, η^2^ = 0.183; HD: atomoxetine vs vehicle- early treatment block: Sidak p = 0.001, d = 0.50; late treatment block: Sidak p = 0.012, d = 0.38] while **F)** having no effect on drinking in any of the other groups [all other groups- atomoxetine vs vehicle- early treatment block: all Sidak ps ≥ 0.117; late block: all Sidak ps≥0.350]. Data are presented as group means ± SEM, with data from individual subjects shown in **A**,**C**&**E**. Blue shaded regions indicate baseline data ie. The last three drug-free sessions prior to the commencement of treatment. **\***p<0.05 Sidak-corrected group comparison for the indicated groups.

After twenty-one SIP sessions, at a time the control over coping behaviour has devolved to the aDLS dopamine-dependent habit system (13), HD, IHD, ILD and LD rats were each randomly assigned to receive vehicle or atomoxetine administration (1.0mg/kg, i.p.) prior to each daily SIP session for 17 days, with groups matched for baseline water consumption (**Figure 5A,C&E**). Chronic atomoxetine administration selectively exacerbated hyperdipsia, as revealed by a substantial increase in the level of hyperdipsia in HD rats, compared to vehicle-treated HD controls (**Figure 5D**), with no effect on adjunctive drinking in LD (**Figure 5D**), or on SIP in IHD or ILD (**Figure 5F**) rats.

### Molecular correlates of the exacerbation of compulsive coping by atomoxetine

#### Immediate early gene (IEG) hotspot analysis

Chronic atomoxetine treatment increased cFos mRNA levels in both the aDLS and the pDMS (**Figure 6A**), while it decreased zif268 mRNA levels in the former (**Figure 6B**) in all rats. No differences between compulsive and non-compulsive rats (see **SOM**) and no other effects of atomoxetine were found in any other region of interest (**Figure 6C&D**).

**Figure 6.**
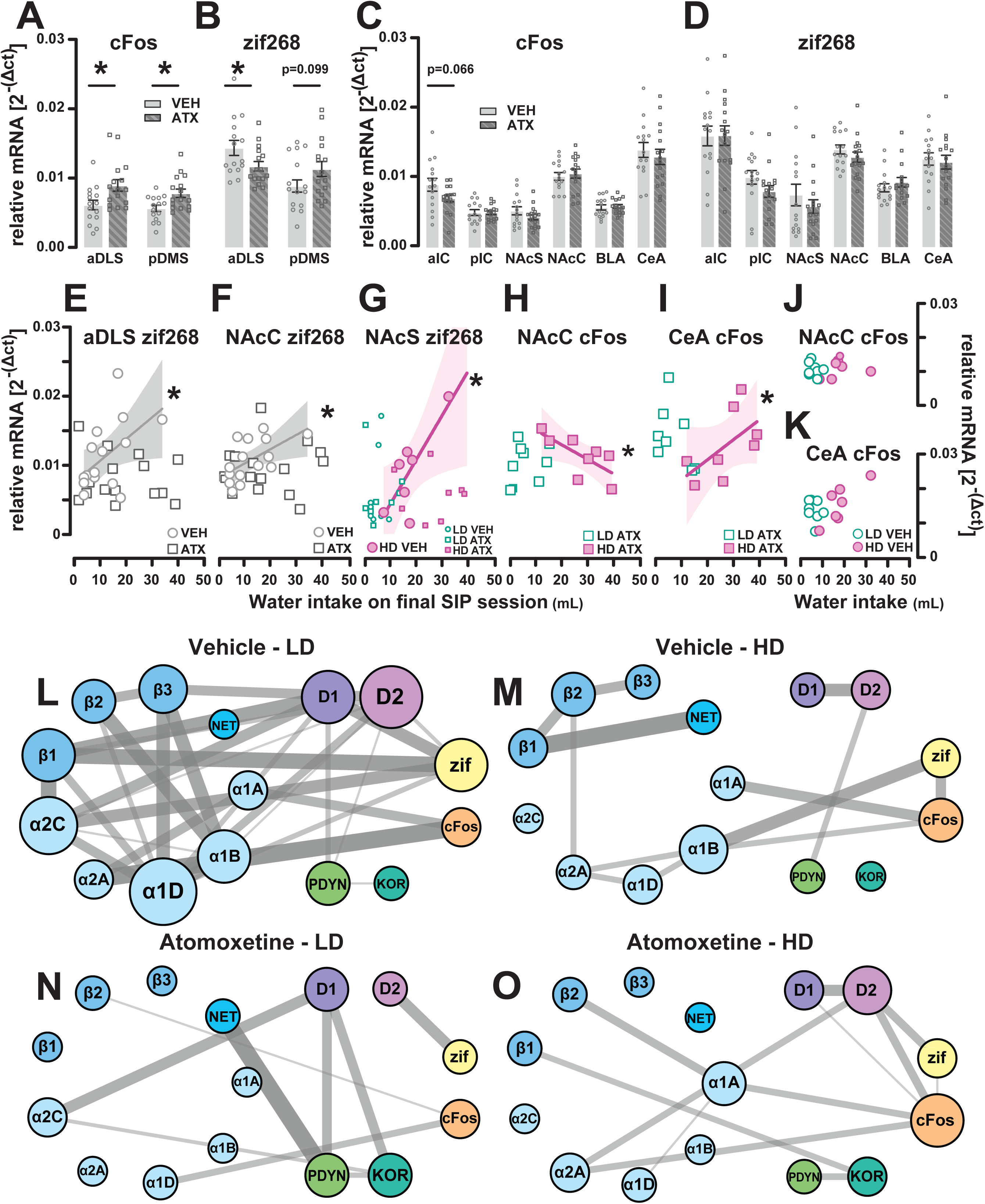
Chronic atomoxetine alters SIP-related cellular activity and plasticity within compulsion-relevant brain regions and results in unique transcriptional signatures within the NacS of compulsive high-drinking and non-polydipsic low-drinking rats. **A)** Compared to vehicle controls, atomoxetine-treated rats exhibited greater mRNA levels of the activity-related marker cFos in both the aDLS [treatment: F_1,27_ = 6.881, p = 0.014, η^2^ = 0.203] and pDMS [treatment: F_1,27_ = 4.913, p = 0.035, η^2^=0.154], and **B)** a concomitant reduction in expression of the plasticity marker zif268 selectively in the aDLS [treatment: F_1,27_ = 5.347, p = 0.029, η^2^ = 0.165]. **C-D)** IEG levels in the insula cortex, nucleus accumbens, and amygdala were comparable between treatment groups [Fs ≤ 3.668, ps ≥ 0.066]. High and low drinkers did not differ in IEG expression across all regions of interest (see **SOM**). In vehicle-treated rats, zif268 expression in the **E)** aDLS [rho = 0.524, p = 0.045] and **F)** NAcC [rho = 0.596, p = 0.019] positively correlated with levels of SIP. **G)** NAcS zif268 levels positively correlated with water drinking, yet only in vehicle-treated compulsive HD rats [rho = 0.788, p = 0.035; all other groups: ps ≥ 0.643]. Selectively in atomoxetine-treated HDs, cFos levels in the **H)** NAcC negatively related to SIP drinking [rho = -0.762, p = 0.028], while **I)** CeA cFos levels positively correlated with water consumption [rho = 0.690, p = 0.058]. **J, K)** No significant cFos-behaviour correlations were observed in vehicle-treated HDs, nor LDs [ps ≥ 0.094]. In the NAcS, expression of IEGs and various target genes of the noradrenaline, dopamine, endogenous opioid systems were differentially correlated in **L)** vehicle-treated rats that failed to acquire polydipsia (LD) versus **M)** those which developed hyperdipsia (HD) [see **SOM** for a table of all correlation coefficients and p-values]. Notably, **M)** HD controls exhibited fewer overall correlations between target genes than **L)** vehicle-treated LDs. Chronic atomoxetine treatment also resulted in distinct transcriptional patterns in **N)** LD and **O)** HD rats. For **K-N**), node size represents the number of significant correlations a given gene has with other target genes, as an index of its weight in the transcriptional network. Edge width and opacity are proportional to the Spearman rho value between two nodes, and only correlations which remained significant after Benjamini-Hochberg correction are displayed. Node colour indicates the gene category: blue = noradrenaline (receptors and transporter), green = dopamine receptors, purple = kappa endogenous opioid system (peptide and receptor), yellow = IEG, orange = plasticity-related IEG. **\*and line**: p < 0.05 Sidak-corrected comparison. **\* alone:** p < 0.059 Spearman rank correlation.

In vehicle-treated rats, irrespective of adjunctive drinking group, the level of water intake was associated with zif268 mRNA levels in the aDLS and the NAcC (**Figure 6E&F**), while levels of hyperdipsia in vehicle-treated HD rats were specifically associated with zif268 mRNA levels in the NAcS (**Figure 6G**). This relationship was absent in atomoxetine-treated HDs rats, which instead demonstrated a negative correlation between hyperdipsia and cFos mRNA levels in the NAcC alongside a positive relationship between hyperdipsia and cFos mRNA levels in the CeA that were not present in their vehicle-treated counterparts, respectively (**Figure 6H&I**). Neither NAcC or CeA cFos levels significantly correlated with water consumption in LDs nor in vehicle-treated HDs (**Figure 6J&K**), and no other significant correlations were observed.

#### Effect of chronic atomoxetine on the compulsion-related NAcS transcriptional landscape

Within the NAcS, where zif268 mRNA was selectively related to hyperdipsia (**Figure 6G**), the mRNA levels of each gene of interest did not differ between vehicle- or atomoxetine-treated HD and LD rats (**Supplementary Figure 1A-B**). In vehicle-treated HD rats, mRNA levels of the noradrenergic receptors (NArs) did not tend to correlate with cFos and zif268 (**Figure 6M**), in marked contrast to the pattern observed in LD rats (**Figure 6L**). Indeed, in vehicle-treated LD rats, cFos and zif268 mRNA levels were closely related to those of several NArs, in a system where all NAr mRNA levels were highly correlated to each other (**Figure 6L**). These compulsion-like-specific NAcS transcriptional relationships were profoundly altered by atomoxetine (**Figure 6N-O**). In LD rats, atomoxetine resulted in a loss of many NAr-NAr and NAr-IEG correlations, accompanied by emergent kappa/dynorphin-D1 dopamine receptor and D2-Dar-zif268 mRNA correlations (**Figure 6N**). Atomoxetine-treated HD rats showed emergent correlations between DArs and IEGs mRNA levels at the expense of those observed between all three β NAr subtypes in their vehicle-treated counterparts (**Figure 6O**). All Spearman rho values are presented in **Supplementary Table 2**.

## Discussion

The present results reveal that, at the population level, sign-tracking facilitates the acquisition of coping behaviour manifested as adjunctive drinking and confers increased vulnerability to develop its compulsive manifestation, hyperdipsia (11, 13), while goal-tracking protects against this transition. Additionally, goal-tracking predicts the magnitude of SIP in a subgroup of rats that adopt an adaptive coping strategy.

In contrast to the protective effect that chronic (but not acute) atomoxetine treatment has against the development of hyperdipsia in vulnerable highly impulsive rats (26), here it selectively exacerbated hyperdipsia once behaviour was well-established and mediated by the aDLS dopamine-dependent habit system (13). This suggests that while atomoxetine may be a safe, or even protective, medication for ADHD patients who have not yet engaged in maladaptive coping strategies or have a low compulsivity trait (23), it may instead worsen compulsivity or precipitate compulsion in patients exhibiting nascent compulsive-like behaviour, as suggested in two case studies (53, 54).

At the neural systems level, atomoxetine abolished the compulsion-like-specific relationship between drinking behaviour and activity-related plasticity in the NAcS, and engaged cellular activity across the dorsal striatum. Atomoxetine decreased mRNA levels of the plasticity marker zif268 (55), indicating a reduction in activity-related plasticity in the aDLS. Chronic atomoxetine also disrupted the pattern of SIP-related plasticity in the incentive habit-related NAcC-aDLS circuit (48), the activation of which was associated with the magnitude of adjunctive drinking in vehicle controls, regardless of its adaptive or compulsive nature. These are intriguing effects, considering that only the NAcS receives noradrenergic innervation (11, 51, 52, 56).

Within the NAcS, zif268 mRNA levels were uniquely related to the severity of hyperdipsia. An exploratory transcriptional network analysis indicated that compulsive HD rats exhibited less transcriptional convergence within the NAcS (i.e., fewer correlations) between mRNA levels of IEGs and candidate genes of the catecholamine and endogenous opioid systems, than LD rats. In the former, chronic atomoxetine treatment further decreased intra-NAcS correlations between mRNA levels of noradrenaline receptors and IEGs in favour of emergent correlations between DA receptor and IEG mRNA levels. This suggests a switch from a noradrenergic to a dopaminergic dominance in the catecholaminergic influence on coping-related NAcS cellular activity and plasticity.

Both sign-tracking and the acquisition of SIP depend on dopaminergic mechanisms in the NAc (21, 36, 57). These overlapping dopaminergic mechanisms may explain the facilitated development of SIP observed in ST rats. Interestingly, while sign-tracking confers an increased vulnerability to develop a high SIP or hyperdipsic phenotype, as determined categorically with a quartile split classification, the difference in adjunctive drinking between ST and GT rats progressively decreased when SIP had become well-established and habitual (13). The difference in coping behaviour between STs and GTs may be accounted for by their differential behavioural response to Pavlovian cues (30, 32, 34). One hypothesis for future research is that the SIP chamber itself may act as an occasion setter for the negative reinforcing properties of the drinking-induced anxiolysis that accompanies the development of SIP (13, 15), which may be differentially imbued with incentive salience by STs and GTs (58).

Goal-tracking was instead associated with the tendency to develop low levels of SIP in intermediate SIP rats. In these ILD rats, goal-tracking correlated with the magnitude of the adjunctive strategy, a relationship not found in hyperdipsic, compulsive HD nor in non-polydipsic LD rats. Thus, goal-tracking may predict the predilection toward moderate and adaptive coping strategies. This is in line with recent evidence showing that stronger goal-tracking tendencies relate to reduced alcohol consumption in rats (59). Further research will be necessary to investigate whether the similar representativity of GT and ST rats in the LD group represents an equal contribution of each trait to the tendency to rely on alcohol to cope with stress, which characterises LD individuals (13, 18, 60).

Goal-tracking was reduced specifically in GTs by acute atomoxetine. This effect is only partially in agreement with the previous demonstration that acute atomoxetine influences both sign- and goal-tracking (46). Several methodological differences may explain these discrepancies, such as the much younger age at which animals in the present study were characterised (70 vs ∼100 days, respectively), or the way in which GT and ST groups were defined. The previous study used a tripartite Pavlovian conditioned approach index to classify STs and GTs (46) rather than a clustering algorithm applied to the sign- and goal-tracking behaviours (CS-lever contacts and CS-magazine entries). Together with a tendency for younger individuals to display higher sign- tracking tendencies (61), the data-driven classification approach in the present study may result in the identification of a smaller population of ST rats (22% vs 40%) characterised by a more extreme sign-tracking phenotype (∼90 CS-lever contacts per session vs ∼65) which is potentially less pervious to acute atomoxetine. The dissociation between the lack of sensitivity of sign-tracking to atomoxetine and the noradrenergic basis of both impulsivity and the development of hyperdipsia suggests a dual neurochemical basis of the vulnerability to compulsive coping behaviourally manifested as impulsivity and sign-tracking.

Acute atomoxetine also had no effect on well-established SIP, thereby suggesting that a chronic treatment, such as used in Experiment 2, and the consequent neuroplastic adaptations, are necessary to influence habitual, aDLS DA-dependent compulsive coping behaviour.

Indeed, chronic atomoxetine treatment exacerbated well-established hyperdipsia in HD rats, opposite to its effect on the development of hyperdipsia in vulnerable highly impulsive rats under the exact same conditions (26). Taken together, these results demonstrate that neuroplastic mechanisms engaged by protracted engagement in coping behaviour are associated with profound alterations in noradrenergic signalling. This aligns with evidence that the development of hyperdipsia is accompanied by a downregulation of the plasticity-related gene Arc in a subset of LC neurons in HD rats (11). Therefore, while non-vulnerable rats may be able to engage adaptive plasticity mechanisms during the development of ventral striatum dopamine-dependent SIP to maintain control over their coping behaviour, vulnerable HD rats cannot do it without potentiation of noradrenergic signalling by atomoxetine. The aberrant plasticity within noradrenergic systems that emerges with protracted coping in HD rats (11) results in a diametrically opposite control over behaviour between the acquisition of adjunctive coping and its manifestation as an aDLS dopamine-dependent compulsive habit (13). Further research is warranted to elucidate the precise mechanisms of this shift. However, the distinct effects of repeated atomoxetine on the development of ventral striatum dopamine-dependent SIP and the expression of aDLS dopamine-dependent hyperdipsia demonstrates the key role of noradrenergic mechanisms in the control of the latter via at least two corticostriatal circuits involved in compulsive coping behaviour (23).

Chronic exposure to atomoxetine can induce enduring functional changes via neuroplastic alterations within the ventral striatal catecholamine system (62) and other corticolimbic networks (63, 64). At the cellular level, the effects of chronic atomoxetine treatment can be mediated by activation of pre-synaptic α2-autoreceptors, which would result in a reduction in spontaneous LC neuronal firing and noradrenaline release (65) or alterations in the expression or transduction of specific postsynaptic noradrenergic and other monoaminergic receptors in response to chronically elevated extracellular NA levels (62). Coupled with dysregulated plasticity within the noradrenergic LC of compulsive rats (11), such adaptations in the NAcS may contribute to the exacerbation of hyperdipsia by atomoxetine through remote control over aDLS dopamine-dependent mechanisms underlying compulsive behaviours (Belin et al., 2008 ; 66, 67-69) via the ascending dopamine-dependent spiralling circuitry (70).

The NAcS transcription patterns explored in the present study start to shed light on such compulsion-specific adaptations in the mesolimbic system and their alteration by atomoxetine. Compared to vehicle-treated LD rats in which mRNA levels of every NAr and DAr subtype were correlated with at least two other catecholamine receptors, vehicle-treated HDs exhibited very little convergence in the level of expression of NArs and DArs, suggesting a compulsion-specific divergence in the regulation of post-synaptic monoaminergic receptors. Additionally, vehicle- treated HDs also showed few correlations between NArs and DArs and the two markers of cellular activity and plasticity, cFos and zif268. This is in marked contrast to the very high convergence of mRNA levels of both IEGS and those of NArs and DArs observed in the NAcS of vehicle-treated LDs, whereby five catecholamine markers correlated with zif268 and two NArs correlated with cFos. Together, this indicates an uncoupling of NAr-related mechanisms from those associated with cellular activity and plasticity in the NAcS in high drinkers, as opposed to the shared involvement of noradrenergic and dopaminergic mechanisms observed in LD rats. Such compulsion-related alterations are consistent with the evidence for the independent, but interacting, role the two catecholamines play in compulsive coping and compulsion vulnerability traits like impulsivity (11, 41–43, 71) .

Interestingly, atomoxetine resulted in a similar disruption of correlations between NAr and IEG mRNA levels in the NAcS in all rats, demonstrating the ability of chronic noradrenergic pharmacological manipulation to influence the relationship between NAr expression in and the functional engagement (activation and activity-related plasticity) of the NAcS (72, 73). In compulsive rats, atomoxetine-exacerbated hyperdipsia was accompanied by emerging correlations between the mRNA levels of DArs and those of cFos and zif268, suggesting a drug- induced functional coupling between dopaminergic mechanisms and those mediating the functional engagement of the NAcS. One may hypothesise that this reflects a treatment-induced re-engagement of the DA-dependent NAc activity which otherwise underlies SIP acquisition (21, 74). At a time where coping responding is dependent on the aDLS dopamine-dependent habit system, such putative reengagement of ventral striatal dopamine-dependent motivational processes may contribute to an increased vigor of the compulsive habit (48). In this context, given that dopaminergic mechanisms in the ventral striatum influence the functional engagement of both the pDMS and the aDLS (75, 76), the atomoxetine-induced increase in cFos across both dorsal striatal territories observed is not surprising. Indeed, while neither dorsal striatal territory receives noradrenergic innervation (11), they are both under functional control of catecholaminergic mechanisms in the NAcS via spiralling striato-meso-striatal connections (70). In addition, aDLS DA-dependent maladaptive habits are also maintained by functional interactions with the CeA (47), which was shown here to be recruited by atomoxetine in HD rats, as revealed by the emergent correlation between the level of hyperdipsia and CeA cFos expression. Atomoxetine treatment also blunted SIP-related plasticity (as assessed by zif268 mRNA levels) within the incentive habit system (namely NAcC-aDLS) (48), which otherwise tracked adjunctive drinking in vehicle-treated controls. Together these data suggest that chronic atomoxetine treatment aberrantly engaged dopamine-dependent motivational mechanisms within a rigid incentive habit system, originating in the ventral striatum, thereby exacerbating compulsion in HD rats.

It is important to acknowledge the exploratory and descriptive nature of the transcriptional network analysis in Experiment 2. Thus, the interpretations derived from these transcriptomic analyses offer insight into the overall transcriptional signature of atomoxetine-invigorated compulsive-like behaviour and provide a theoretical framework from which future mechanistic experiments can be developed. These experiments should also investigate whether these effects are observed in females. Nevertheless, these data reveal novel relationships between sign-tracking and hyperdipsia vs goal-tracking and controlled coping, and further elucidate the differential contribution, and associated transcriptional signature, of noradrenergic mechanisms in the development vs the expression of compulsive coping, mediated by the ventral and dorsolateral striatum, respectively. These data also provide novel insights into the mechanisms by which atomoxetine induces or worsens compulsions in humans (53, 54).

## Supporting information

Supplemental Methods, Results, and Tables

## Data availability statement

The data can be available at: link to Cambridge repository will be added upon acceptance.

## Authors contributions

DB and ABR designed the experiments. ABR, AHM, ST, BN, CY and MP carried out the behavioural experiments. ABR and MF carried out the molecular biology experiments. CC, SJ and DB analysed and interpreted the data and wrote the manuscript.

## Acknowledgements and funding

This work, carried out at the Department of Psychology of the University of Cambridge, was funded by UKRI grants to DB (MR/W019647/1 and (MR/N02530X/1 (PI: Barry Everitt), a Leverhulme Trust Early Career (ECF-2018-713) and Isaac Newton Trust fellowship (18.08(g)) to MF and Cambridge Trust International Scholarship (10712081-2023) and NSERC Postgraduate Scholarship – Doctoral (PGSD3-579465–2023) to CC. The authors would like to thank the technical staff of the University Biomedical Services of the Combined Facility for the husbandry and care provided to the animals involved in this research. A preprint of this work was available on bioRxiv: https://www.biorxiv.org/content/10.1101/2024.10.08.617254v1.

## References

1. Cabib S, Campus P, Colelli V. Learning to cope with stress: psychobiological mechanisms of stress resilience. Rev Neurosci. 2012;23(5-6):659–72.

2. McLafferty M, Armour C, Bunting B, Ennis E, Lapsley C, Murray E, et al. Coping, stress, and negative childhood experiences: The link to psychopathology, self-harm, and suicidal behavior. Psych J. 2019;8(3):293–306.

3. Moritz S, Jahns AK, Schroder J, Berger T, Lincoln TM, Klein JP, et al. More adaptive versus less maladaptive coping: What is more predictive of symptom severity? Development of a new scale to investigate coping profiles across different psychopathological syndromes. J Affect Disord. 2016;191:300–7.

4. Matheson K, Anisman H. Systems of coping associated with dysphoria, anxiety and depressive illness: a multivariate profile perspective. Stress. 2003;6(3):223–34.

5. Wood RT, Griffiths MD. A qualitative investigation of problem gambling as an escape-based coping strategy. Psychol Psychother. 2007;80(Pt 1):107–25.

6. Goodwin H, Haycraft E, Meyer C. Emotion regulation styles as longitudinal predictors of compulsive exercise: a twelve month prospective study. J Adolesc. 2014;37(8):1399–404.

7. Holt LJ, Armeli S, Tennen H, Austad CS, Raskin SA, Fallahi CR, et al. A person-centered approach to understanding negative reinforcement drinking among first year college students. Addict Behav. 2013;38(12):2937–44.

8. Falk JL. The nature and determinants of adjunctive behavior. Physiol Behav. 1971;6(5):577–88.

9. Falk J. Production of polydipsia in normal rats by an intermittent food schedule. Science. 1961;133(3447):195-6.

10. Dantzer R, Terlouw C, Mormede P, Le Moal M. Schedule-induced polydipsia experience decreases plasma corticosterone levels but increases plasma prolactin levels. Physiol Behav. 1988;43(3):275–9.

11. Velazquez-Sanchez C, Muresan L, Marti-Prats L, Belin D. The development of compulsive coping behaviour is associated with a downregulation of Arc in a Locus Coeruleus neuronal ensemble. Neuropsychopharmacology. 2023;48(4):653–63.

12. Levine S, Weinberg J, Brett LP. Inhibition of pituitary-adrenal activity as a consequence of consummatory behavior. Psychoneuroendocrinology. 1979;4(4):275–86.

13. Marti-Prats L, Giuliano C, Domi A, Puaud M, Pena-Oliver Y, Fouyssac M, et al. The development of compulsive coping behavior depends on dorsolateral striatum dopamine-dependent mechanisms. Mol Psychiatry. 2023;28(11):4666–78.

14. Belin-Rauscent A, Daniel ML, Puaud M, Jupp B, Sawiak S, Howett D, et al. From impulses to maladaptive actions: the insula is a neurobiological gate for the development of compulsive behavior. Mol Psychiatry. 2016;21(4):491–9.

15. Tazi A, Dantzer R, Mormede P, Le Moal M. Pituitary-adrenal correlates of schedule-induced polydipsia and wheel running in rats. Behav Brain Res. 1986;19(3):249–56.

16. Merchan A, Sanchez-Kuhn A, Prados-Pardo A, Gago B, Sanchez-Santed F, Moreno M, et al. Behavioral and biological markers for predicting compulsive-like drinking in schedule-induced polydipsia. Prog Neuropsychopharmacol Biol Psychiatry. 2019;93:149–60.

17. Pellon R, Ruiz A, Moreno M, Claro F, Ambrosio E, Flores P. Individual differences in schedule-induced polydipsia: neuroanatomical dopamine divergences. Behav Brain Res. 2011;217(1):195–201.

18. Fouyssac M, Puaud M, Ducret E, Marti-Prats L, Vanhille N, Ansquer S, et al. Environment-dependent behavioral traits and experiential factors shape addiction vulnerability. Eur J Neurosci. 2021;53(6):1794–808.

19. Moreno M, Flores P. Schedule-induced polydipsia as a model of compulsive behavior: neuropharmacological and neuroendocrine bases. Psychopharmacology (Berl). 2012;219(2):647–59.

20. Flores P, Sánchez-Kuhn A, Merchán A, Vilches O, García-Martín S, Moreno M. Schedule-Induced Polydipsia: Searching for the Endophenotype of Compulsive Behavior. World Journal of Neuroscience. 2014;04(03):253–60.

21. Robbins TW, Koob GF. Selective disruption of displacement behaviour by lesions of the mesolimbic dopamine system. Nature. 1980;285(5764):409-12.

22. Everitt BJ, Robbins TW. Drug Addiction: Updating Actions to Habits to Compulsions Ten Years On. Annu Rev Psychol. 2016;67:23–50.

23. Robbins TW, Banca P, Belin D. From compulsivity to compulsion: the neural basis of compulsive disorders. Nat Rev Neurosci. 2024;25(5):313–33.

24. Belin D, Mar AC, Dalley JW, Robbins TW, Everitt BJ. High Impulsivity Predicts the Switch to Compulsive Cocaine-Taking. Science. 2008;320(5881):1352-5.

25. Jones JA, Belin-Rauscent A, Jupp B, Fouyssac M, Sawiak SJ, Zuhlsdorff K, et al. Neurobehavioral Precursors of Compulsive Cocaine Seeking in Dual Frontostriatal Circuits. Biological Psychiatry Global Open Science. 2023.

26. Ansquer S, Belin-Rauscent A, Dugast E, Duran T, Benatru I, Mar AC, et al. Atomoxetine decreases vulnerability to develop compulsivity in high impulsive rats. Biol Psychiatry. 2014;75(10):825–32.

27. Lovic V, Saunders BT, Yager LM, Robinson TE. Rats prone to attribute incentive salience to reward cues are also prone to impulsive action. Behav Brain Res. 2011;223(2):255–61.

28. Schettino M, Ceccarelli I, Tarvainen M, Martelli M, Orsini C, Ottaviani C. From skinner box to daily life: Sign-tracker phenotype co-segregates with impulsivity, compulsivity, and addiction tendencies in humans. Cognitive, Affective, & Behavioral Neuroscience. 2022;22(6):1358–69.

29. Swintosky M, Brennan JT, Koziel C, Paulus JP, Morrison SE. Sign tracking predicts suboptimal behavior in a rodent gambling task. Psychopharmacology (Berl). 2021;238(9):2645–60.

30. Lashley RL, Rosellini RA. Modulation of schedule-induced polydipsia by Pavlovian conditioned states. Physiol Behav. 1980;24(2):411–4.

31. Plonsky M, Driscoll CD, Warren DA, Rosellini RA. Do random time schedules induce polydipsia in the rat? Animal Learning & Behavior. 1984;12(4):355–62.

32. Hamm RJ, Porter JH, Kaempf GL. Stimulus generalization of schedule-induced polydipsia. J Exp Anal Behav. 1981;36(1):93–9.

33. Shurtleff D, Delamater AR, Riley AL. A reevaluation of the CS− hypothesis for schedule- induced polydipsia under intermittent schedules of pellet delivery. Animal Learning & Behavior. 1983;11(2):247–54.

34. Porter JH, Hamm RJ. Associative control of schedule-induced drinking. Animal Learning & Behavior. 1984;12(3):339–40.

35. Flagel SB, Clark JJ, Robinson TE, Mayo L, Czuj A, Willuhn I, et al. A selective role for dopamine in stimulus-reward learning. Nature. 2011;469(7328):53-7.

36. Fraser KM, Janak PH. Long-lasting contribution of dopamine in the nucleus accumbens core, but not dorsal lateral striatum, to sign-tracking. Eur J Neurosci. 2017;46(4):2047–55.

37. Economidou D, Theobald DE, Robbins TW, Everitt BJ, Dalley JW. Norepinephrine and dopamine modulate impulsivity on the five-choice serial reaction time task through opponent actions in the shell and core sub-regions of the nucleus accumbens. Neuropsychopharmacology. 2012;37(9):2057–66.

38. Benn A, Robinson ES. Differential roles for cortical versus sub-cortical noradrenaline and modulation of impulsivity in the rat. Psychopharmacology (Berl). 2017;234(2):255–66.

39. Weissenborn R, Blaha CD, Winn P, Phillips AG. Schedule-induced polydipsia and the nucleus accumbens: electrochemical measurements of dopamine efflux and effects of excitotoxic lesions in the core. Behav Brain Res. 1996;75(1-2):147–58.

40. Moreno M, Gutierrez-Ferre VE, Ruedas L, Campa L, Sunol C, Flores P. Poor inhibitory control and neurochemical differences in high compulsive drinker rats selected by schedule- induced polydipsia. Psychopharmacology (Berl). 2012;219(2):661–72.

41. Moreno M, Cardona D, Gomez MJ, Sanchez-Santed F, Tobena A, Fernandez-Teruel A, et al. Impulsivity characterization in the Roman high- and low-avoidance rat strains: behavioral and neurochemical differences. Neuropsychopharmacology. 2010;35(5):1198–208.

42. de Villiers AS, Russell VA, Sagvolden T, Searson A, Jaffer A, Taljaard JJ. Alpha 2- adrenoceptor mediated inhibition of [3H]dopamine release from nucleus accumbens slices and monoamine levels in a rat model for attention-deficit hyperactivity disorder. Neurochem Res. 1995;20(4):427–33.

43. Russell V, Allie S, Wiggins T. Increased noradrenergic activity in prefrontal cortex slices of an animal model for attention-deficit hyperactivity disorder--the spontaneously hypertensive rat. Behav Brain Res. 2000;117(1-2):69–74.

44. Vanhille N, Belin-Rauscent A, Mar AC, Ducret E, Belin D. High locomotor reactivity to novelty is associated with an increased propensity to choose saccharin over cocaine: new insights into the vulnerability to addiction. Neuropsychopharmacology. 2015;40(3):577–89.

45. Chernoff CS, Hynes TJ, Winstanley CA. Noradrenergic contributions to cue-driven risk- taking and impulsivity. Psychopharmacology (Berl). 2021;238(7):1765–79.

46. Holden JM. Effects of two kinds of noradrenergic ADHD medicines on sign-tracking and goal-tracking in male rats. Exp Clin Psychopharmacol. 2022;30(6):760–73.

47. Murray JE, Belin-Rauscent A, Simon M, Giuliano C, Benoit-Marand M, Everitt BJ, et al. Basolateral and central amygdala differentially recruit and maintain dorsolateral striatum- dependent cocaine-seeking habits. Nat Commun. 2015;6:10088.

48. Belin D, Belin-Rauscent A, Murray JE, Everitt BJ. Addiction: failure of control over maladaptive incentive habits. Curr Opin Neurobiol. 2013;23(4):564–72.

49. Marti-Prats L, Belin-Rauscent A, Fouyssac M, Puaud M, Cocker PJ, Everitt BJ, et al. Baclofen decreases compulsive alcohol drinking in rats characterized by reduced levels of GAT- 3 in the central amygdala. Addict Biol. 2021;26(4):e13011.

50. Magnard R, Fouyssac M, Vachez YM, Cheng Y, Dufourd T, Carcenac C, et al. Pramipexole restores behavioral inhibition in highly impulsive rats through a paradoxical modulation of frontostriatal networks. Transl Psychiatry. 2024;14(1):86.

51. Delfs JM, Zhu Y, Druhan JP, Aston-Jones GS. Origin of noradrenergic afferents to the shell subregion of the nucleus accumbens: anterograde and retrograde tract-tracing studies in the rat. Brain Res. 1998;806(2):127–40.

52. Berridge CW, Stratford TL, Foote SL, Kelley AE. Distribution of dopamine beta- hydroxylase-like immunoreactive fibers within the shell subregion of the nucleus accumbens. Synapse. 1997;27(3):230–41.

53. Araz Altay M. Atomoxetine Induced Obsessive-Compulsive Disorder. Hospital Practices and Research. 2019;4(2):68–70.

54. İnce C, Karakuş M, Karadeniz S, Kandil S. Turkish Journal of Pediatric Disease. 2015;9(2):140–2.

55. Knapska E, Kaczmarek L. A gene for neuronal plasticity in the mammalian brain: Zif268/Egr-1/NGFI-A/Krox-24/TIS8/ZENK? Prog Neurobiol. 2004;74(4):183–211.

56. Schroeter S, Apparsundaram S, Wiley RG, Miner LH, Sesack SR, Blakely RD. Immunolocalization of the cocaine- and antidepressant-sensitive l-norepinephrine transporter. J Comp Neurol. 2000;420(2):211–32.

57. Saunders BT, Robinson TE. The role of dopamine in the accumbens core in the expression of Pavlovian-conditioned responses. Eur J Neurosci. 2012;36(4):2521–32.

58. Fraser KM, Janak PH. Occasion setters attain incentive motivational value: implications for contextual influences on reward-seeking. Learn Mem. 2019;26(8):291–8.

59. Hakus A, Foo JC, Casquero-Veiga M, Gül AZ, Hintz F, Rivalan M, et al. Sex-associated differences in incentive salience and drinking behaviour in a rodent model of alcohol relapse. Addiction Biology. 2025;30(1).

60. Marti-Prats L, Belin D. Characterization in the rat of the individual tendency to rely on alcohol to cope with distress and the ensuing vulnerability to drink compulsively. Brain Commun. 2024;6(3):fcae169.

61. DeAngeli NE, Miller SB, Meyer HC, Bucci DJ. Increased sign-tracking behavior in adolescent rats. Dev Psychobiol. 2017;59(7):840–7.

62. Ruocco LA, Carnevale UA, Treno C, Sadile AG, Melisi D, Arra C, et al. Prepuberal subchronic methylphenidate and atomoxetine induce different long-term effects on adult behaviour and forebrain dopamine, norepinephrine and serotonin in Naples high-excitability rats. Behav Brain Res. 2010;210(1):99–106.

63. Fumagalli F, Cattaneo A, Caffino L, Ibba M, Racagni G, Carboni E, et al. Sub-chronic exposure to atomoxetine up-regulates BDNF expression and signalling in the brain of adolescent spontaneously hypertensive rats: comparison with methylphenidate. Pharmacol Res. 2010;62(6):523–9.

64. Sun H, Cocker PJ, Zeeb FD, Winstanley CA. Chronic atomoxetine treatment during adolescence decreases impulsive choice, but not impulsive action, in adult rats and alters markers of synaptic plasticity in the orbitofrontal cortex. Psychopharmacology (Berl). 2012;219(2):285–301.

65. Bari A, Aston-Jones G. Atomoxetine modulates spontaneous and sensory-evoked discharge of locus coeruleus noradrenergic neurons. Neuropharmacology. 2013;64:53–64.

66. Belin D, Everitt BJ. Cocaine seeking habits depend upon dopamine-dependent serial connectivity linking the ventral with the dorsal striatum. Neuron. 2008;57(3):432–41.

67. Belin D, Everitt BJ. The Neural and Psychological Basis of a Compulsive Incentive Habit. In: Steiner H, tseng K, editors. Handbook of basal ganglia structure and function. Handbook of Behavioral Neuroscience: Elsevier, ACADEMIC PRESS; 2010. P. 571-92.

68. Belin D, Economidou D, Pelloux Y, Everitt BJ. Habit Formation and Compulsion. Animal Models of Drug Addiction. 2011;53:337–78.

69. Haber SN, Fudge JL, McFarland NR. Striatonigrostriatal pathways in primates form an ascending spiral from the shell to the dorsolateral striatum. J Neurosci. 2000;20(6):2369–82.

70. Ikemoto S. Dopamine reward circuitry: two projection systems from the ventral midbrain to the nucleus accumbens-olfactory tubercle complex. Brain Res Rev. 2007;56(1):27–78.

71. Russell VA. The nucleus accumbens motor-limbic interface of the spontaneously hypertensive rat as studied in vitro by the superfusion slice technique. Neurosci Biobehav Rev. 2000;24(1):133–6.

72. Jones MW, Errington ML, French PJ, Fine A, Bliss TV, Garel S, et al. A requirement for the immediate early gene Zif268 in the expression of late LTP and long-term memories. Nat Neurosci. 2001;4(3):289–96.

73. Filipkowski RK, Knapska E, Kaczmarek L. c-Fos and Zif268 in Learning and Memory— Studies on Expression and Function. In: Pinaud R, Tremere LA, editors. Immediate Early Genes in Sensory Processing, Cognitive Performance and Neurological Disorders. Boston, MA: Springer US; 2006. P. 137-58.

74. Mittleman G, Whishaw IQ, Jones GH, Koch M, Robbins TW. Cortical, hippocampal, and striatal mediation of schedule-induced behaviors. Behav Neurosci. 1990;104(3):399–409.

75. O’Doherty J, Dayan P, Schultz J, Deichmann R, Friston K, Dolan RJ. Dissociable roles of ventral and dorsal striatum in instrumental conditioning. Science. 2004;304(5669):452-4.

76. Yin HH, Ostlund SB, Balleine BW. Reward-guided learning beyond dopamine in the nucleus accumbens: the integrative functions of cortico-basal ganglia networks. Eur J Neurosci. 2008;28(8):1437–48.

