## Supplemental Methods, Results, and Tables for "Compulsive coping behaviour, developed predominantly by sign-trackers, is exacerbated by chronic atomoxetine"

Chernoff *et al.*

### Supplementary Methods

#### Subjects

One hundred and eight 10-week-old adult male Sprague-Dawley rats (Charles River, UK) weighing between 280-355g at the start of the experiment were single-housed under a reversed 12h light/dark cycle (lights off at 7:00 AM) in temperature and humidity-controlled rooms. After a week of habituation to the animal facility, they were food restricted to 85-90% or 80-83% of theoretical free-feeding weight prior to autoshaping or SIP training, respectively. Water was freely available in the homecage. One rat in Experiment 2 was excluded due to excessive weight loss, despite intensive supportive care. Another rat in Experiment 2 was excluded because it played with the bottle spout, emptying the entire 300mL SIP water bottle without necessarily drinking. All procedures were conducted under the project license 70/8072 held by David Belin in accordance with the requirements of the UK Animals (Scientific Procedures) Act 1986, amendment 2012.

#### Apparatus

Behavioural testing occurred in standard operant chambers (24x25.4x26.7cm) (Med Associates, St. Albans, VT) located within ventilated sound-attenuating cupboards. Autoshaping chambers were outfitted with a house light, a food magazine through which dustless food pellets (BioServ, USA) could be delivered, two retractable metal levers flanking the food magazine, and two stimulus lights located on the wall above each lever. SIP chambers were equipped with a house light, a food magazine, and a receptacle in which a water bottle with a stainless-steel sipper spout could be installed, which was situated opposite to the food magazine. Infrared beams were installed in the food magazines and on either side of the water bottle spout to record magazine head entries and licks, respectively. Chambers were controlled by MedPC IV software (Med Associates Inc., Ltd) using code written by ABR, MF, and MP.

#### Behavioural training

##### Autoshaping

The autoshaping task was conducted in twelve dedicated MedAssociates operant boxes.

###### Habituation and magazine training

To habituate rats to the food pellets used as reward during autoshaping and SIP procedures, each rat was given free access to 20 pellets in the homecage one day before beginning operant training. The following day, rats underwent a magazine training session, during which 50 pellets were delivered into the food magazine following a variable interval 30 seconds schedule.

###### Autoshaping training

For the following five days, rats were trained in an autoshaping, pavlovian conditioned approach task, which involved 25 presentations of the lever and cue light above [ie. the compound conditioned stimulus (CS)] for 8 seconds, after which the lever was retracted, the stimulus light turned off, and one food pellet was delivered into the magazine. CS-US pairs were presented according to a 90-second variable time schedule. Sign-tracking was quantified by the number of lever presses made during the CS presentation, as an index of approach toward and interaction with either the light or the lever of the CS. Goal-tracking was measured by the number of nose pokes made into the food magazine during the CS presentation

##### Schedule-induced polydipsia (SIP)

SIP training was carried out in twenty-four dedicated MedAssociates operant boxes as described previously (1).

###### Habituation and baseline water consumption

Rats in Experiment 2 first underwent habituation to the food pellets through free access to 20 pellets in the homecage, the day prior to being habituated to the operant chamber. Rats in Experiment 1, which were previously habituated to the food pellets prior to autoshaping training, went straight on to chamber habituation. During operant habituation, rats were placed into the chambers (31.8 x 25.4 x 26.7 cm) for 60 minutes with free access to a water bottle and 60 immediately available food pellets, placed in the magazine. The water bottle was situated on the wall opposite to the food magazine. Water consumption during the 60-minute chamber habituation session was recorded as a measure of baseline water consumption [pre-session water bottle weight (g) – post-session water bottle weight (g)].

###### Magazine training

Following habituation, rats performed one 60-minute session of magazine training, which involved the delivery of 60 food pellets on a random time 60-second schedule into the food magazine. Water consumption throughout the 60-minute session was recorded. The next day SIP training began.

###### SIP training

Rats underwent 18 (Experiment 1) or 21 (Experiment 2) manipulation-free SIP sessions, over the course of which individual differences in polydipsic coping responses emerged. During each 60-minute SIP session, rats had free access to a 300 mL water bottle, and food pellets were delivered into the food magazine on a fixed time 60 second (FT60s) schedule. The house light was illuminated throughout the entirety of the session. Head entries into the food magazine and licks at the waterspout were automatically recorded by infrared beams. Water consumption was recorded [pre-session water bottle weight (g) – post-session water bottle weight (g)] as the main metric of schedule-induced polydipsia.

For every dosing day, the SIP session was identical to those performed during training. Rats were exposed to the SIP procedure 6 days a week starting at the same time of day for each rat throughout the entire experiment.

#### Atomoxetine treatment

For the acute atomoxetine challenge in Experiment 1, each rat received intraperitoneal (i.p) administration of 0, 1.0, and 3.0mg/kg at 1 mL/kg, counterbalanced across days and rats according to a Latin Square design, and following a repeating 3-day schedule: 1) drug-free behaviour session – no injection, 2) dosing session – pre-session injection, and 3) rest – no behavioural testing or injection. In Experiment 2, following 21 days of drug-free SIP acquisition, rats were randomly assigned to receive atomoxetine (1.0 mg/kg at 1.0 mL/kg i.p (2)) or vehicle (1 mL/kg 0.9% saline, i.p) prior to each SIP session for the next 17 days. Within each drinking group (ie. HD, IHD, ILD, and LD), randomly assigned treatment groups were matched for baseline pre-treatment water consumption. All injections occurred 30 minutes before behavioural testing.

#### Molecular biology analyses - Quantitative polymerase chain reaction (qPCR)

##### Tissue collection

Forty-five minutes following the final SIP session, rats were deeply anaesthetized with isoflurane and decapitated. Brains were rapidly extracted and flash frozen in -40°C isopentane for ~3 minutes before being stored at -80°C. Brains were then coronally sliced at 300 µm on a cryostat, and regions of interest [ie. anterior (aIC) and posterior insula (pIC), nucleus accumbens shell (NAcS) and core (NAcC), anterior dorsolateral striatum (aDLS), posterior dorsomedial striatum (pDMS), and basolateral (BLA) and central amygdala (CeA)] were identified based on the Rat Brain Atlas (3) and bilaterally micro-dissected using a 1mm micro-puncher.

##### RNA extraction and reverse transcription

RNA was extracted from brain tissue micro-punches using the Quick-RNA Microprep Kit (Zymo Research) following manufacturer guidelines, and RNA levels were quantified using a NanoDrop ND-1000 spectrophotometer (Thermo Fisher Scientific). cDNA banks were reverse-transcribed from RNA using the RT^2^ First Strand Kit (Qiagen, UK) following the manufacturer’s guidelines.

##### Quantification and analysis

For every brain region of interest, mRNA levels for the immediate early genes (IEGs) cFos and zif268 were quantified. Samples taken from the NAcS were also processed to assess the mRNA levels corresponding to various noradrenaline and dopamine receptors, the noradrenaline transporter, the kappa opioid receptor, and prodynorphin, as well as those of both cFos and zif268. Primers used to assess the mRNA levels for the various target genes are listed in **Supplementary Table 1**. Real-time PCR (RT-PCR) was performed using the CFx96 Real-Time PCR Detection System (Bio-Rad, UK), with RT^2^ SYBR Green Mastermix (Qiagen, UK) to quantify amplified DNA. RT-PCR was conducted on samples of 25 µL (1µL cDNA + 24 µL PCR reaction mix), beginning with 1 initial step of 95°C for 10 mins, followed by 40 cycles of 95°C for 15 seconds plus 60°C for 60 seconds. Relative mRNA levels of each target gene were calculated using CFX Manager Software (Bio-Rad, UK) and expressed as 2^-∆CT^ relative to the housekeeping gene, cyclophilin A. Each sample was run in duplicate for every brain region and target mRNA, with the average relative expression of the two runs used for the final relative mRNA level.

#### Data and statistical analyses

Assumptions of normality, homogeneity of variance, and sphericity were tested using Shapiro-Wilk, Levene, and Mauchley tests, respectively. In instances where normality was violated, as for some relative mRNA levels (**Supplementary Table 2**), data were log-transformed before analysis. If sphericity was violated, the Huynh-Feldt correction was applied. The relative distribution of subjects across SIP and autoshaping groups were compared using a Chi-squared test and z-test of independent proportions. Behavioural data were subjected to Analyses of Variance (ANOVA) with group (ST vs GT, or HD, IHD, ILD, LD) and/or treatment (vehicle vs atomoxetine) as between-subject factors, and time (daily session) as the within-subject factor.

On confirmation of significant main effects or interactions, differences among individual means were further analysed using Sidak-corrected post-hoc pairwise comparisons. Effect sizes for ANOVA effects and pairwise comparisons were reported as partial eta squared (η^2^) and Cohen’s d, respectively. Multivariate and simple linear regression was used to assess the extent to which variation in performance on the autoshaping task or SIP could predict behaviour on the other paradigm. For the hotspot analysis, univariate ANOVA with SIP group and treatment as between-subject factors was used to compare IEG mRNA levels between groups for the compulsion-relevant brain regions listed above. Relationships between behavioural and molecular variables were examined using Spearman correlations. Transcription networks containing gene-gene Spearman correlations which survived Benjamini-Hochberg correction were used to visually explore and describe transcription patterns within the NAcS of vehicle- or atomoxetine-treated HD and LD rats.

Figures were generated using GraphPad Prism 10.2 (GraphPad, USA) and Adobe Illustrator (Adobe, CA, USA), and network plots were created using Cytoscape 3.10 (Institute for Systems Biology, USA).

##### Behavioural classification

Baseline behavioural performance in both the autoshaping and SIP procedures was determined based on an individual’s average level of responding (ie. CS lever presses and CS magazine entries for the former and water consumption for the latter) across the last three pre-treatment sessions in either paradigm.

For sign/goal tracking, rats were identified as sign-trackers (STs), intermediate performers (INT), or goal-trackers (GTs) using a K-means cluster analysis of CS contacts and magazine entries across the last three autoshaping sessions. Such a clustering approach allows an unbiased objective identification of subpopulations characterized by both their sign and goal-tracking responses. The k-means algorithm was instructed to form three clusters consistent with the three groups commonly defined in other studies of individual differences in autoshaping (ie. ST, INT, GT) (2, 3).

For SIP, rats from both experiments were stratified by a quartile split of baseline SIP water consumption and classified as high (HD), high intermediate (IHD), low intermediate (ILD), and low (LD) drinkers. This approach segregates hyperdipsic rats (HD) from rats which do not develop an adjunctive coping response (LD) and maximizes the distance between these two groups (2,6–9), while reducing type I and preventing type II errors (4). Further, rats which develop adaptive low levels of adjunctive coping (ILD) and those which acquire polydipsia, yet do not exhibit extreme, compulsive hyperdipsia (IHD), form the intermediate two quartiles of the population flanking the median. These intermediate groups, which develop different degrees of controlled polydipsic drinking, have hitherto seldom been considered in studies investigating the mechanisms of hyperdipsia (5), often being amalgamated with the HD and LD groups (6, 7). This introduces type II errors and occludes our understanding of the mechanisms underlying adaptive polydipsic behaviour. Thus, including these subgroups provides a richer understanding of individual differences and their associated biobehavioural mechanisms across the whole spectrum of coping behaviour.

The relative distribution of STs and GTs across the four SIP groups was assessed using a Chi-square test, followed by a z-test for independent proportions.

##### Behavioural performance and drug challenge

Longitudinal SIP and autoshaping data were analyzed using repeated measures analysis of variance (ANOVA) with session as a within-subject factor. SIP sessions were divided into equal-sized blocks of training and treatment sessions (early/late training or treatment, where applicable) to assess potential group differences across distinct epochs of the experiment. For all repeated measures analyses of SIP data, block was included as a within-subjects factor. To compare task performance between groups in Experiment 1, autoshaping cluster (GT/ST) and drinking group (HD/IHD/ILD/LD) were set as between-subject factors. Acute atomoxetine challenge data were analyzed using repeated measures ANOVA with dose (3 levels: 0,1,3mg/kg) as a within-subject factor, and autoshaping group (ST/GT) and drinking group (HD/IHD/ILD/LD) as between-subject factors. To investigate the effect of chronic atomoxetine treatment in Experiment 2, and how this may differ between rats which develop distinct coping strategies, treatment (atomoxetine/vehicle) and drinking group (HD/IHD/ILD/LD) were included as between-subject factors. Any significant main effects or interactions were further examined by Sidak-corrected pairwise comparisons between appropriate group means.

Linear regression of baseline performance was conducted as follows, for each SIP group separately, to assess the predictive relationship between autoshaping and SIP, and whether the relationship differs between drinking groups. Multivariate linear regression, with both CS lever presses and CS magazine entries as predictors, was used to assess the extent to which both autoshaping traits together predict subsequent SIP. Univariate linear regression, with either CS lever presses or CS magazine entries as the predictor and water consumption as the outcome variable, examined whether SIP drinking could be predicted by either autoshaping trait alone.

##### Immediate early gene (IEG) hotspot analysis

To identify neural “hotspots” specific to adaptive vs compulsive coping behaviour, the relative mRNA levels of the IEGs cFos and zif268 were assessed in the eight brain regions of interest using univariate ANOVA with treatment (atomoxetine/vehicle) and compulsivity group (HD/LD) as between-subject factors. IEG levels in each region of interest were also correlated with water consumption during the final SIP session, using Spearman correlation to further investigate the relationship between regional activation patterns and polydipsic coping behaviour.

##### NAcS transcriptional network analysis

To explore the influence of atomoxetine on the compulsive coping-related transcriptional landscape of the NAcS, the only striatal territory receiving noradrenergic inputs (6,12,13), the mRNA levels of several genes implicated in noradrenaline and dopamine transmission, i.e., noradrenaline and dopamine receptors and the noradrenaline transporter (NET) (14–21), as well as negative affective states, e.g. the dynorphin/kappa opiate receptor system (22–24); were assessed alongside those of the IEGs c-fos and zif268 (25–28) (**Supplementary Table 1**). The levels of each mRNA were first compared between groups using a univariate ANOVA with treatment (ATO vs veh) and group (HD/LD) as between-subject factors. To explore the influence of atomoxetine on the compulsive coping-specific transcriptional landscape, relationships between mRNA levels of each candidate gene were assessed using Spearman correlations were performed for each treatment and SIP group (HD/LD) separately. Resultant p-values were adjusted using the Benjamini-Hochberg correction for multiple comparisons. Only correlations that survived correction were visualized using network plots, whereby each node represented a gene, and edge thickness/opacity corresponded to the correlation’s Spearman Rho value. Non-significant correlations were omitted, and node size was proportional to the number of significant correlations a given target had with other genes.

Supplementary Table 1. List of primers used in the qPCR experiments*.*

All primers were run on samples from the NAcS, and primers for cFos (Fos) and zif268 (Egr1) immediate early genes were used on samples from every brain region of interest (see methods).

| Target gene | Symbol | UniGene no. | Qiagen  catalogue no. | Experiment |
| --- | --- | --- | --- | --- |
| Adrenergic receptor, α1A | Adra1a | Rn.9991 | PPR06850A | NAcS network |
| Adrenergic receptor, α1B | Adra1b | Rn.10032 | PPR06754A | NAcS network |
| Adrenergic receptor, α1D | Adra1d | Rn.11314 | PPR06783A | NAcS network |
| Adrenergic receptor, α2A | Adra2a | Rn.170171 | PPR06801B | NAcS network |
| Adrenergic receptor, α2C | Adra2c | Rn.10297 | PPR06768A | NAcS network |
| Adrenergic receptor, β1 | Adrb1 | Rn.87064 | PPR06825A | NAcS network |
| Adrenergic receptor, β2 | Adrb2 | Rn.10206 | PPR06764A | NAcS network |
| Adrenergic receptor, β3 | Adrb3 | Rn.10100 | PPR06759A | NAcS network |
| Noradrenaline transporter (NET) | Slc6a2 | Rn.14577 | PPR06785A | NAcS network |
| Dopamine receptor, D1A | Drd1 | Rn.24039 | PPR06790A | NAcS network |
| Dopamine receptor, D2 | Drd2 | Rn.87299 | PPR06827A | NAcS network |
| Opioid receptor, kappa 1 (KOR) | Oprk1 | Rn.89571 | PPR06833G | NAcS network |
| Prodynorphin (PDYN) | Pdyn | Rn.44471 | PPR49782A | NAcS network |
| FBJ osteosarcoma oncogene (cFos) | Fos | Rn.103750 | PPR55248C | Network & hotspot |
| Early growth response (Zif268, zif) | Egr1 | Rn.9096 | PPR44272B | Network & hotspot |
| Cyclophilin A (housekeeping gene) | Ppia | Rn.1463 | PPR06504A | All |

Supplementary Table 2. List of relative mRNA levels that were subjected to log-transformation in order to be analysed using parametric analyses*.*

All genes the relative mRNA level of which was log transformed prior to analysis, due to violations of normality as per the Shapiro-Wilk test, are indicated with a check mark. Genes that were normally distributed were not log-transformed, and analyses were carried out on raw, untransformed data.

| Target gene | Brain region | Log-transformed |
| --- | --- | --- |
| Adrenergic receptor, α1A | NAcS | **✓** |
| Adrenergic receptor, α1B | NAcS | **✓** |
| Adrenergic receptor, α1D | NAcS | **✓** |
| Adrenergic receptor, α2A | NAcS |  |
| Adrenergic receptor, α2C | NAcS |  |
| Adrenergic receptor, β1 | NAcS |  |
| Adrenergic receptor, β2 | NAcS |  |
| Adrenergic receptor, β3 | NAcS | **✓** |
| Noradrenaline transporter (NET) | NAcS | **✓** |
| Dopamine receptor, D1A | NAcS | **✓** |
| Dopamine receptor, D2 | NAcS | **✓** |
| Opioid receptor, kappa 1 (KOR) | NAcS |  |
| Prodynorphin (PDYN) | NAcS |  |
| FBJ osteosarcoma oncogene (cFos) | aIC |  |
|  | pIC |  |
|  | NAcS |  |
|  | NAcC |  |
|  | BLA | **✓** |
|  | CeA |  |
|  | aDLS | **✓** |
|  | pDMS |  |
| Early growth response (Zif268, zif) | aIC |  |
|  | pIC |  |
|  | NAcS | **✓** |
|  | NAcC |  |
|  | BLA | **✓** |
|  | CeA |  |
|  | aDLS | **✓** |
|  | pDMS |  |

### Supplementary Results

Emergence of individual differences over the course of autoshaping and SIP training

##### Experiment 1

Both CS lever pressing (ie. sign-tracking) and CS magazine entries (ie. goal-tracking) behaviours were acquired across the five autoshaping training sessions [CS lever presses- session: F_4,180_ = 33.161, p < 0.001, η^2^ = 0.424; session 1 vs 5: Sidak p < 0.001, d = 1.176; CS magazine entries- session: F_4,180_ = 41.071, p < 0.001, η^2^ = 0.483; session 1 vs 5: Sidak p < 0.001, d = 1.334]. As expected, STs and GTs demonstrated greater responding either on the lever or at the food magazine, respectively, during the CS presentation [CS lever presses- session x group: F_8,180_ = 17.240, p < 0.001, η^2^ = 0.434; GTs vs STs- session 1: Sidak p = 0.003, d = 0.504; all other sessions: Sidak ps < 0.001; CS magazine entries- session x group: F_8,180_ = 23.766, p < 0.001, η^2^ = 0.514; GTs vs STs- session 1: Sidak p = 0.094; session 2: Sidak p < 0.001, d = 1.165; all other sessions: Sidak ps < 0.001]**.**

Adjunctive water drinking was acquired by the third day of SIP training at the population level (main effect of session: F_17,784_ = 55.614, p < 0.001, η^2^ = 0.558; session 1 vs 2: Sidak p = 0.238, session 1 vs 3: Sidak p < 0.001, d = 1.221; session 1 vs all other sessions: all ps ≤ 0.001). Differences between SIP groups emerged at various points in acquisition, such that by the end of SIP training, groups exhibited distinct levels of water consumption across the last three SIP training sessions (BL) [session x SIP group: F_51,782_ = 3.402, p < 0.001, η^2^ = 0.188; HD vs LD- sessions 1-2: Sidak ps ≥ 0.770; sessions 3-18: Sidak ps ≤ 0.042, ds ≥ 0.408; HD vs ILD- sessions 1-5: Sidak ps ≥ 0.087; sessions 6-18: Sidak ps ≤ 0.036, ds ≥ 0.416; HD vs IHD- sessions 1-7: Sidak ps ≥ 0.188; sessions 8-18: Sidak ps ≤ 0.012, ds ≥ 0.475; IHD vs LD- sessions 1-11: Sidak ps ≥ 0.085; sessions 12-18: Sidak ps ≤ 0.003, ds ≥ 0.545; IHD vs ILD- sessions 1-14: Sidak ps ≥ 0.152; sessions 15-18 ps ≤ 0.037, ds ≥ 0.393; ILD v LD- sessions 1-12: Sidak ps ≥ 0.534; sessions 13-18: Sidak ps ≤ 0.054, ds ≥ 0.393]**.**

##### Experiment 2

As in Experiment 1, distinct phenotypes based on adjunctive water drinking emerged across 21 days of SIP training [session x SIP group: F_60,980_ = 4.952, p < 0.001, η^2^ = 0.408], whereby compulsive HD rats differed from non-polydipsic LD rats as early as the second session [HD vs LD- sessions 1: p = 0.066; sessions 2-21: ps ≤ 0.054, ds ≥ 0.392]. HDs drank significantly more water than the low (ILD) and high (IHD) intermediate drinking groups from session 11 and 18, respectively [HD vs ILD- sessions 1-10: p = 0.081; sessions 11-21: ps ≤ 0.029, ds ≥ 0.380; HD vs IHD- sessions 1-17: p = 0.173; sessions 18-21: ps ≤ 0.043, ds ≥ 0.300]. The intermediate drinking groups diverged from session 12 onward [IHD vs ILD- sessions 1-11: Sidak ps ≥ 0.138; sessions 12-21: ps ≤ 0.044, ds ≥ 0.403], and non-polydipsic LDs showed less water drinking than either ILD or IHD polydipsic groups from session 20 and 9, respectively [ILD vs LD- sessions 1-19: Sidak ps ≥ 0.140; sessions 20-21: ps ≤ 0.034, ds ≥ 0.382; IHD vs LD- sessions 1-8: p = 0.073; sessions 9-21: ps ≤ 0.007, ds ≥ 0.445].

Following SIP acquisition, rats were randomly assigned to receive daily pre-session atomoxetine or vehicle injections throughout the ensuing treatment period. Atomoxetine and vehicle treatment groups were matched for pre-treatment water consumption within each group [treatment group: F_1,50_ = 0.432, p = 0.519; treatment group x drinking group: F_3,50_ = 0.754, p = 0.525; atomoxetine vs vehicle: all groups Sidak ps ≥ 0.131].

Supplementary Table 3. Spearman rho values of correlations between mRNA levels of candidate genes in the NAcS*.*

Separate tables display relative mRNA level correlations for vehicle (VEH)-treated LD **A)** and HD **B)** rats, and atomoxetine (ATX)-treated LD **C)** and HD **D)** rats. Bold values indicate correlations that remained significant following Benjamini-Hochberg correction.

| A. VEH - LD | 1 | 2 | 3 | 4 | 5 | 6 | 7 | 8 | 9 | 10 | 11 | 12 | 13 | 14 | 15 |
| --- | --- | --- | --- | --- | --- | --- | --- | --- | --- | --- | --- | --- | --- | --- | --- |
| 1. α1A | -- |  |  |  |  |  |  |  |  |  |  |  |  |  |  |
| 2. α1B | 0.60 | -- |  |  |  |  |  |  |  |  |  |  |  |  |  |
| 3. α1D | 0.55 | **0.93** | -- |  |  |  |  |  |  |  |  |  |  |  |  |
| 4. α2A | **0.88** | 0.55 | 0.48 | -- |  |  |  |  |  |  |  |  |  |  |  |
| 5. α2C | 0.69 | **0.76** | **0.88** | 0.50 | -- |  |  |  |  |  |  |  |  |  |  |
| 6. β1 | 0.67 | 0.71 | **0.86** | 0.55 | **0.98** | -- |  |  |  |  |  |  |  |  |  |
| 7. β2 | 0.60 | **0.95** | **0.81** | 0.50 | 0.64 | 0.57 | -- |  |  |  |  |  |  |  |  |
| 8. β3 | 0.83 | **0.94** | **0.94** | 0.77 | 0.77 | 0.71 | **0.89** | -- |  |  |  |  |  |  |  |
| 9. NET | 0.57 | 0.71 | 0.68 | 0.14 | 0.68 | 0.54 | 0.79 | 0.60 | -- |  |  |  |  |  |  |
| 10. D1 | 0.50 | 0.62 | **0.81** | 0.43 | **0.90** | **0.95** | 0.48 | 0.54 | 0.39 | -- |  |  |  |  |  |
| 11. D2 | 0.50 | **0.76** | **0.86** | 0.48 | **0.76** | **0.81** | 0.62 | **0.89** | 0.64 | 0.71 | -- |  |  |  |  |
| 12. KOR | 0.05 | 0.12 | 0.24 | 0.07 | 0.33 | 0.43 | 0.26 | 0.09 | 0.11 | 0.60 | 0.31 | -- |  |  |  |
| 13. PDYN | 0.14 | 0.38 | 0.62 | 0.17 | 0.60 | 0.71 | 0.24 | 0.26 | 0.29 | **0.81** | **0.76** | **0.76** | -- |  |  |
| 14. cFos | **0.88** | 0.55 | 0.48 | **1.00** | 0.50 | 0.55 | 0.50 | 0.77 | 0.14 | 0.43 | 0.48 | 0.07 | 0.17 | -- |  |
| 15. zif | 0.62 | 0.69 | **0.81** | 0.57 | **0.93** | **0.98** | 0.52 | 0.60 | 0.36 | **0.93** | **0.79** | 0.36 | 0.69 | 0.57 | -- |

| B. VEH- HD | 1 | 2 | 3 | 4 | 5 | 6 | 7 | 8 | 9 | 10 | 11 | 12 | 13 | 14 | 15 |
| --- | --- | --- | --- | --- | --- | --- | --- | --- | --- | --- | --- | --- | --- | --- | --- |
| 1. α1A | -- |  |  |  |  |  |  |  |  |  |  |  |  |  |  |
| 2. α1B | 0.75 | -- |  |  |  |  |  |  |  |  |  |  |  |  |  |
| 3. α1D | 0.79 | **0.82** | -- |  |  |  |  |  |  |  |  |  |  |  |  |
| 4. α2A | 0.54 | **0.82** | **0.82** | -- |  |  |  |  |  |  |  |  |  |  |  |
| 5. α2C | 0.04 | 0.32 | 0.21 | 0.57 | -- |  |  |  |  |  |  |  |  |  |  |
| 6. β1 | 0.57 | 0.54 | 0.50 | 0.75 | 0.54 | -- |  |  |  |  |  |  |  |  |  |
| 7. β2 | 0.39 | 0.64 | 0.46 | **0.82** | 0.79 | **0.89** | -- |  |  |  |  |  |  |  |  |
| 8. β3 | 0.03 | 0.60 | 0.37 | 0.83 | 0.71 | 0.54 | **0.89** | -- |  |  |  |  |  |  |  |
| 9. NET | 0.80 | 0.70 | 0.60 | 0.70 | 0.10 | **1.00** | 0.70 | 0.50 | -- |  |  |  |  |  |  |
| 10. D1 | 0.07 | 0.14 | 0.25 | 0.14 | 0.50 | 0.57 | 0.61 | 0.09 | 0.20 | -- |  |  |  |  |  |
| 11. D2 | 0.07 | 0.07 | 0.39 | 0.11 | 0.25 | 0.36 | 0.39 | 0.09 | 0.20 | **0.93** | -- |  |  |  |  |
| 12. KOR | 0.07 | 0.04 | 0.11 | 0.43 | 0.07 | 0.64 | 0.43 | 0.20 | 0.90 | 0.29 | 0.07 | -- |  |  |  |
| 13. PDYN | 0.21 | 0.36 | 0.64 | 0.32 | 0.07 | 0.29 | 0.21 | 0.14 | 0.50 | 0.75 | **0.82** | 0.32 | -- |  |  |
| 14. cFos | **0.89** | **0.82** | 0.64 | 0.64 | 0.14 | 0.75 | 0.64 | 0.26 | 0.90 | 0.43 | 0.39 | 0.29 | 0.07 | -- |  |
| 15. zif | 0.71 | **0.93** | 0.61 | 0.71 | 0.39 | 0.64 | 0.75 | 0.60 | 0.60 | 0.46 | 0.43 | 0.07 | 0.00 | **0.89** | -- |

| C. ATX – LD | 1 | 2 | 3 | 4 | 5 | 6 | 7 | 8 | 9 | 10 | 11 | 12 | 13 | 14 | 15 |
| --- | --- | --- | --- | --- | --- | --- | --- | --- | --- | --- | --- | --- | --- | --- | --- |
| 1. α1A | -- |  |  |  |  |  |  |  |  |  |  |  |  |  |  |
| 2. α1B | 0.38 | -- |  |  |  |  |  |  |  |  |  |  |  |  |  |
| 3. α1D | 0.05 | 0.52 | -- |  |  |  |  |  |  |  |  |  |  |  |  |
| 4. α2A | 0.40 | 0.02 | 0.62 | -- |  |  |  |  |  |  |  |  |  |  |  |
| 5. α2C | 0.24 | 0.00 | 0.24 | 0.10 | -- |  |  |  |  |  |  |  |  |  |  |
| 6. β1 | 0.17 | 0.62 | 0.52 | 0.14 | 0.38 | -- |  |  |  |  |  |  |  |  |  |
| 7. β2 | 0.38 | 0.45 | 0.60 | 0.55 | 0.12 | 0.33 | -- |  |  |  |  |  |  |  |  |
| 8. β3 | 0.14 | 0.77 | 0.49 | 0.20 | 0.31 | 0.43 | 0.31 | -- |  |  |  |  |  |  |  |
| 9. NET | 0.90 | 0.80 | 0.40 | 0.00 | 0.60 | 0.90 | 0.60 | 0.60 | -- |  |  |  |  |  |  |
| 10. D1 | 0.33 | 0.24 | 0.19 | 0.40 | **0.90** | 0.48 | 0.05 | 0.20 | 0.60 | -- |  |  |  |  |  |
| 11. D2 | 0.57 | 0.05 | 0.71 | 0.31 | 0.48 | 0.33 | 0.40 | 0.31 | 0.10 | 0.43 | -- |  |  |  |  |
| 12. KOR | 0.14 | 0.29 | 0.07 | 0.48 | **0.79** | 0.33 | 0.21 | 0.43 | 0.50 | **0.88** | 0.10 | -- |  |  |  |
| 13. PDYN | 0.05 | 0.64 | 0.33 | 0.33 | 0.71 | 0.67 | 0.26 | 0.66 | **1.00** | **0.88** | 0.31 | **0.81** | -- |  |  |
| 14. cFos | 0.05 | 0.43 | **0.83** | 0.45 | 0.17 | 0.19 | **0.76** | 0.54 | 0.40 | 0.17 | 0.71 | 0.05 | 0.26 | -- |  |
| 15. zif | 0.48 | 0.05 | 0.57 | 0.36 | 0.33 | 0.43 | 0.45 | 0.09 | 0.10 | 0.29 | **0.90** | 0.14 | 0.17 | 0.57 | -- |

| D. ATX - LD | 1 | 2 | 3 | 4 | 5 | 6 | 7 | 8 | 9 | 10 | 11 | 12 | 13 | 14 | 15 |
| --- | --- | --- | --- | --- | --- | --- | --- | --- | --- | --- | --- | --- | --- | --- | --- |
| 1. α1A | -- |  |  |  |  |  |  |  |  |  |  |  |  |  |  |
| 2. α1B | 0.62 | -- |  |  |  |  |  |  |  |  |  |  |  |  |  |
| 3. α1D | **0.76** | 0.52 | -- |  |  |  |  |  |  |  |  |  |  |  |  |
| 4. α2A | 0.67 | 0.57 | 0.48 | -- |  |  |  |  |  |  |  |  |  |  |  |
| 5. α2C | 0.38 | 0.24 | 0.19 | 0.43 | -- |  |  |  |  |  |  |  |  |  |  |
| 6. β1 | 0.29 | 0.11 | 0.32 | 0.68 | 0.64 | -- |  |  |  |  |  |  |  |  |  |
| 7. β2 | **0.86** | 0.39 | 0.71 | 0.29 | 0.64 | 0.43 | -- |  |  |  |  |  |  |  |  |
| 8. β3 | 0.90 | 0.90 | 0.90 | 0.70 | 0.50 | 0.20 | 0.80 | -- |  |  |  |  |  |  |  |
| 9. NET | 0.40 | 0.50 | 0.50 | 0.30 | 0.40 | 0.30 | 0.30 | 0.70 | -- |  |  |  |  |  |  |
| 10. D1 | 0.55 | 0.05 | 0.19 | 0.74 | 0.50 | 0.57 | 0.36 | 0.10 | 0.10 | -- |  |  |  |  |  |
| 11. D2 | 0.50 | 0.10 | 0.24 | **0.83** | 0.60 | 0.79 | 0.29 | 0.20 | 0.20 | **0.90** | -- |  |  |  |  |
| 12. KOR | 0.21 | 0.12 | 0.14 | 0.60 | 0.69 | **0.82** | 0.32 | 0.20 | 0.20 | 0.60 | 0.74 | -- |  |  |  |
| 13. PDYN | -0.1 | 0.00 | 0.43 | 0.17 | 0.69 | 0.79 | 0.29 | 0.20 | 0.20 | 0.24 | 0.33 | **0.83** | -- |  |  |
| 14. cFos | **0.83** | 0.43 | 0.52 | **0.83** | 0.64 | 0.64 | 0.68 | 0.70 | 0.70 | **0.76** | **0.86** | 0.62 | 0.19 | -- |  |
| 15. zif | 0.38 | 0.05 | 0.19 | 0.57 | 0.62 | 0.46 | 0.25 | 0.20 | 0.20 | 0.71 | **0.86** | 0.50 | 0.19 | **0.76** | -- |

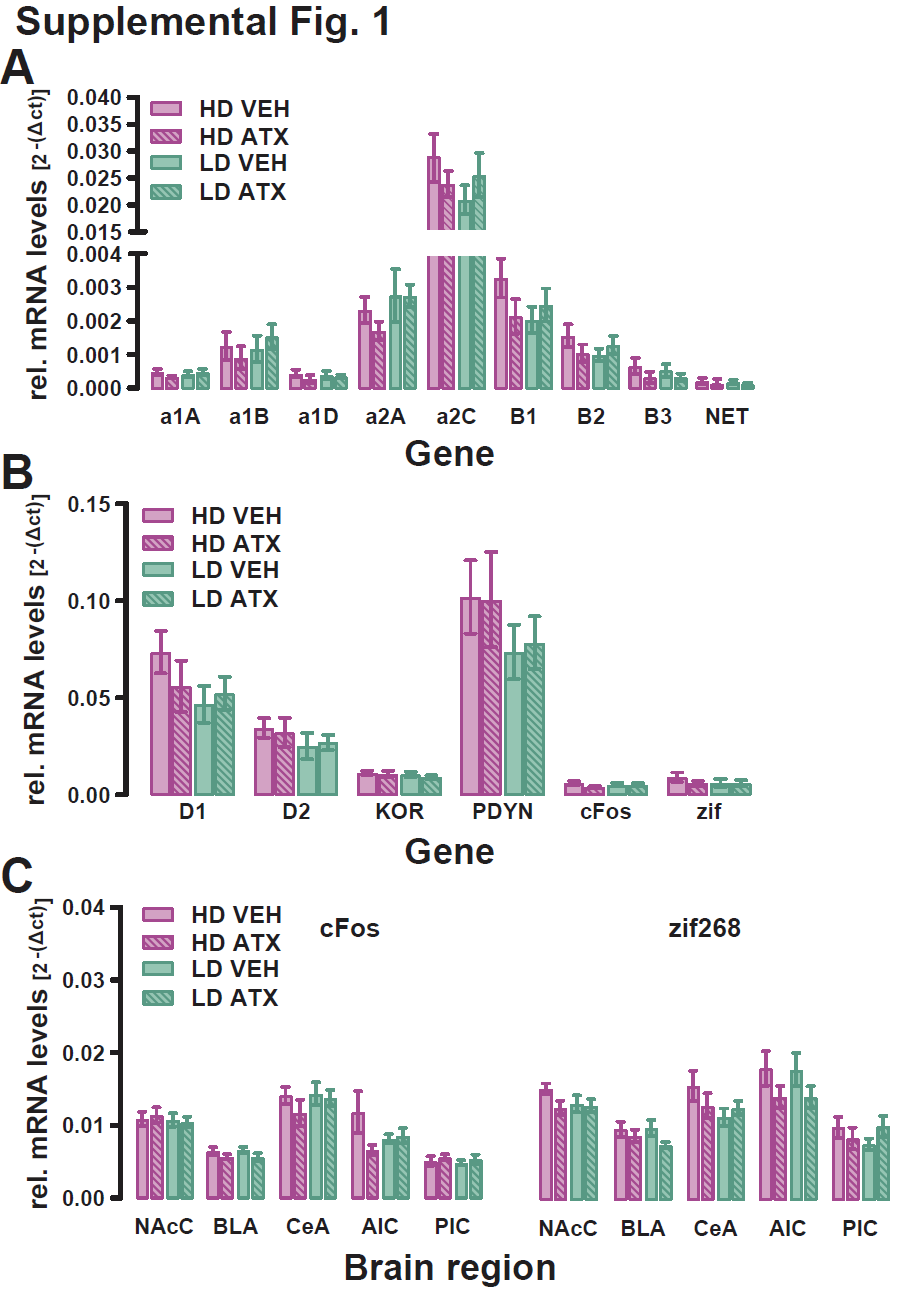

#### Supplementary Figure 1. NAcS mRNA levels of candidate genes were not influenced by coping phenotype or atomoxetine.

The intra-NAcS relative levels of the mRNAs of various genes of interest within the **A)** noradrenaline, **B)** dopamine, and **B)** kappa-prodynorphin endogenous opioid systems, as well as **B)** immediate early genes (IEGs), did not differ between HD and LD rats which received chronic treatment of either atomoxetine or vehicle [all Fs ≤ 2.304; all ps ≥ 0.146]. **C)** IEG expression within all regions of interest also did not differ between treatment and drinking groups [all Fs ≤ 3.668; all ps ≥ 0.066]. Data are presented as mean ± SEM.

### References

1. Ansquer S, Belin-Rauscent A, Dugast E, Duran T, Benatru I, Mar AC, et al. Atomoxetine decreases vulnerability to develop compulsivity in high impulsive rats. Biol Psychiatry. 2014;75(10):825-32.

2. Flagel SB, Watson SJ, Robinson TE, Akil H. Individual differences in the propensity to approach signals vs goals promote different adaptations in the dopamine system of rats. Psychopharmacology (Berl). 2007;191(3):599-607.

3. Jones JA, Belin-Rauscent A, Jupp B, Fouyssac M, Sawiak SJ, Zuhlsdorff K, et al. Neurobehavioral Precursors of Compulsive Cocaine Seeking in Dual Frontostriatal Circuits. Biol Psychiatry Glob Open Sci. 2024;4(1):194-202.

4. Vanhille N, Belin-Rauscent A, Mar AC, Ducret E, Belin D. High locomotor reactivity to novelty is associated with an increased propensity to choose saccharin over cocaine: new insights into the vulnerability to addiction. Neuropsychopharmacology. 2015;40(3):577-89.

5. Fouyssac M, Puaud M, Ducret E, Marti-Prats L, Vanhille N, Ansquer S, et al. Environment-dependent behavioral traits and experiential factors shape addiction vulnerability. Eur J Neurosci. 2021;53(6):1794-808.

6. Pellon R, Ruiz A, Moreno M, Claro F, Ambrosio E, Flores P. Individual differences in schedule-induced polydipsia: neuroanatomical dopamine divergences. Behav Brain Res. 2011;217(1):195-201.

7. Mora S, Merchan A, Aznar S, Flores P, Moreno M. Increased amygdala and decreased hippocampus volume after schedule-induced polydipsia in high drinker compulsive rats. Behav Brain Res. 2020;390:112592.
