## Supplementary figures and images for "Compulsive coping behaviour, developed predominantly by sign-trackers, is exacerbated by chronic atomoxetine"

### Supplementary Figure 1

#### Supplemental Fig. 1

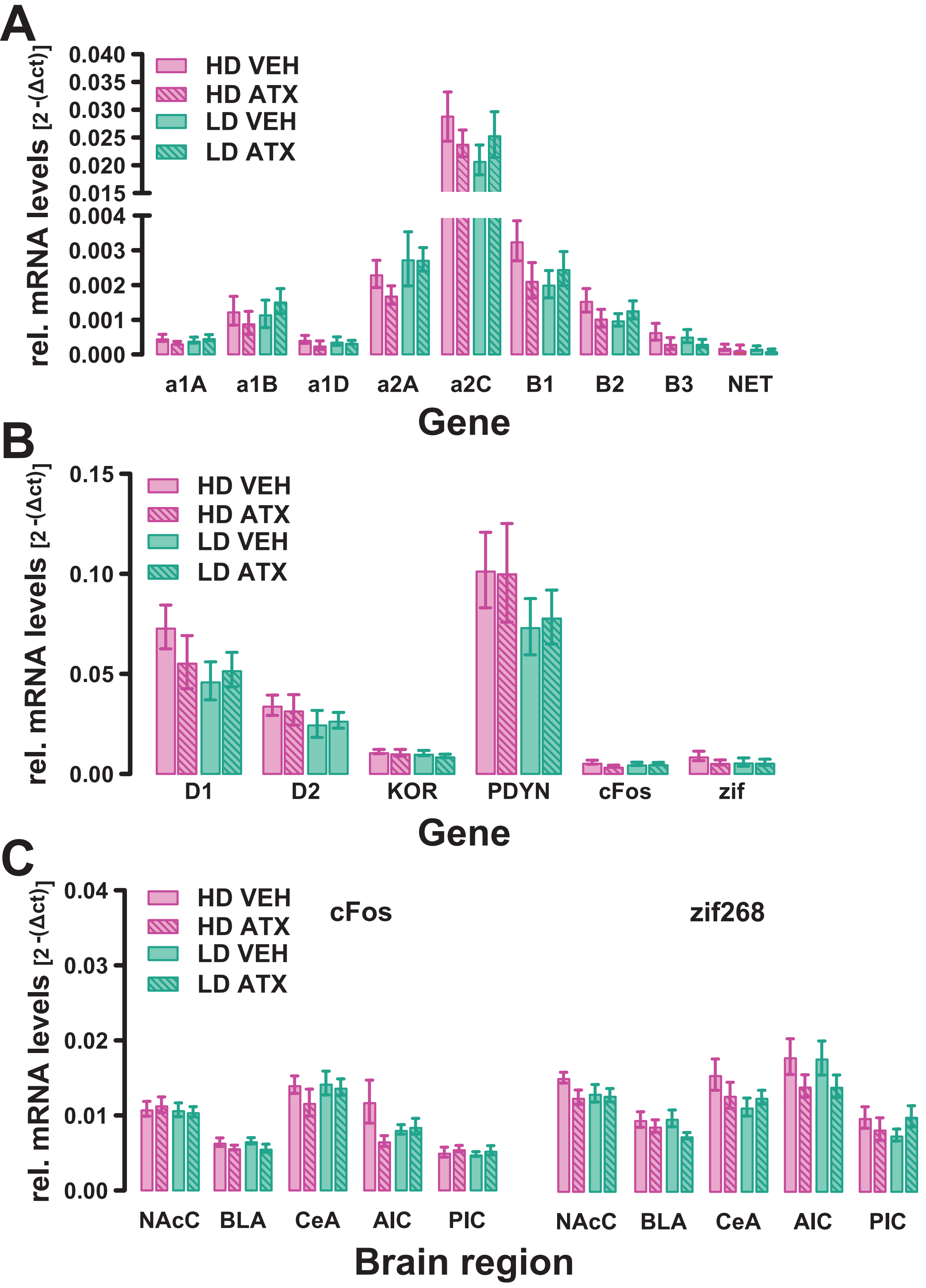
